# An allosteric hot spot in the tandem-SH2 domain of ZAP-70 regulates T-cell signaling

**DOI:** 10.1101/842534

**Authors:** Kaustav Gangopadhyay, Bharat Manna, Swarnendu Roy, Sunitha Kumari, Olivia Debnath, Subhankar Chowdhury, Amit Ghosh, Rahul Das

## Abstract

T-cell receptor (TCR) signaling is initiated by recruiting ZAP-70 to the cytosolic part of TCR. ZAP-70, a non-receptor tyrosine kinase, is composed of an N-terminal tandem SH2 (tSH2) domain connected to the C-terminal kinase domain. The ZAP-70 is recruited to the membrane through binding of tSH2 domain and the doubly-phosphorylated ITAM motifs of CD3 chains in the TCR complex. Our results show that the tSH2 domain undergoes a biphasic structural transition while binding to the doubly-phosphorylated ITAM-ζ1 peptide. The C-terminal SH2 domain binds first to the phosphotyrosine residue of ITAM peptide to form an encounter complex leading to subsequent binding of second phosphotyrosine residue to the N-SH2 domain. We decipher a network of non-covalent interactions that allosterically couple the two SH2 domains during binding to doubly-phosphorylated ITAMs. Mutation in the allosteric network residues, for example, W165C, uncouples the formation of encounter complex to the subsequent ITAM binding thus explaining the altered recruitment of ZAP-70 to the plasma membrane causing autoimmune arthritis in mice. The proposed mechanism of allosteric coupling is unique to ZAP-70, which is fundamentally different from Syk, a close homolog of ZAP-70 expressed in B-cells.

**Significance:** T-cell and B-cell signaling is initiated by the same family of non-receptor tyrosine kinases, ZAP-70 and Syk, respectively. ZAP-70 and Syk share homologous sequence and similar structural architecture, yet the two kinases differ in their mode of ligand recognition. ZAP-70 binds cooperatively to its ligand, whereas Syk binds uncooperatively. Spontaneous mutation (W165C) in the regulatory module of ZAP-70 impairs T-cell signaling causes autoimmune arthritis in SKG mice, the mechanism of which is unknown. We showed that ZAP-70 regulatory module undergoes a biphasic structural transition while binding to its ligand, which is fundamentally different from Syk. We presented a molecular mechanism of cooperativity in ZAP-70 regulatory module that explains altered ligand binding by ZAP-70 mutant found in SKG mice.

## Introduction

The zeta-chain-associated protein tyrosine kinase, ZAP-70, is a non-receptor tyrosine kinase crucial for T-cell signaling, development, activation, and proliferation(1–4). T-cell signaling is commenced by the recruitment of two protein tyrosine kinase, Src family kinase Lck and ZAP-70, to the activated molecular complex of T-cell antigen receptor (TCR)(5, 6). Lck, phosphorylate several tyrosine residues of the immuno-receptor tyrosine-based activation motifs (ITAM) on the intracellular segment of CD3 heterodimer (made up of δ, γ, and ε) and ζ homodimer associated with the TCR(5, 7–10). ZAP-70 is spontaneously recruited to the membrane by binding to the doubly-phosphorylated ITAM (ITAM-Y2P) motifs(11–14). Recruitment of ZAP-70 allows phosphorylation of scaffold proteins that initiates a cascade of downstream biological events(15, 16). The mutation that reduces the ZAP-70 interaction to the ITAM-Y2P motif, for example, W165C mutation in SKG mice attenuate TCR signalling, gives rise to inflammatory arthritis resembling rheumatoid arthritis in human(17).

ZAP-70 has a modular structure comprised of an N-terminal regulatory module connected through a linker (named interdomain-B) to the C-terminal catalytic module (kinase domain)(18) (Figure 1a). The regulatory module is made up of tandem repeats of the Src homology-2 (tSH2) domain connected by a helical linker called interdomain A (Figure 1a). The tSH2 domain has two phosphate-binding pockets, one at the C-terminal SH2 domain (C-SH2) and the second one at the interface of the N-terminal and the C-terminal SH2 domains (N-SH2)(19) (Figure 1b and S2b). In the autoinhibited state, the kinase domain adopts an inactive Cdk/Src-like structure, and the two SH2 domains are separated(20) in an ‘L shaped’ open conformation rendering the tSH2 domain incompatible with binding to ITAM-Y2P-ζ1 peptide(19, 21) (Figure 1b). In the active state, the binding of doubly-phosphorylated ITAM reorient the two SH2 domains with respect to each other in a ‘Y shaped’ close conformation(19, 21) (Figure 1b), facilitate ZAP-70 to take an open conformation(22) resulting in autophosphorylation of regulatory tyrosine residues at the interdomain B and activation loop, respectively (20, 23–29).

**Figure 1:**
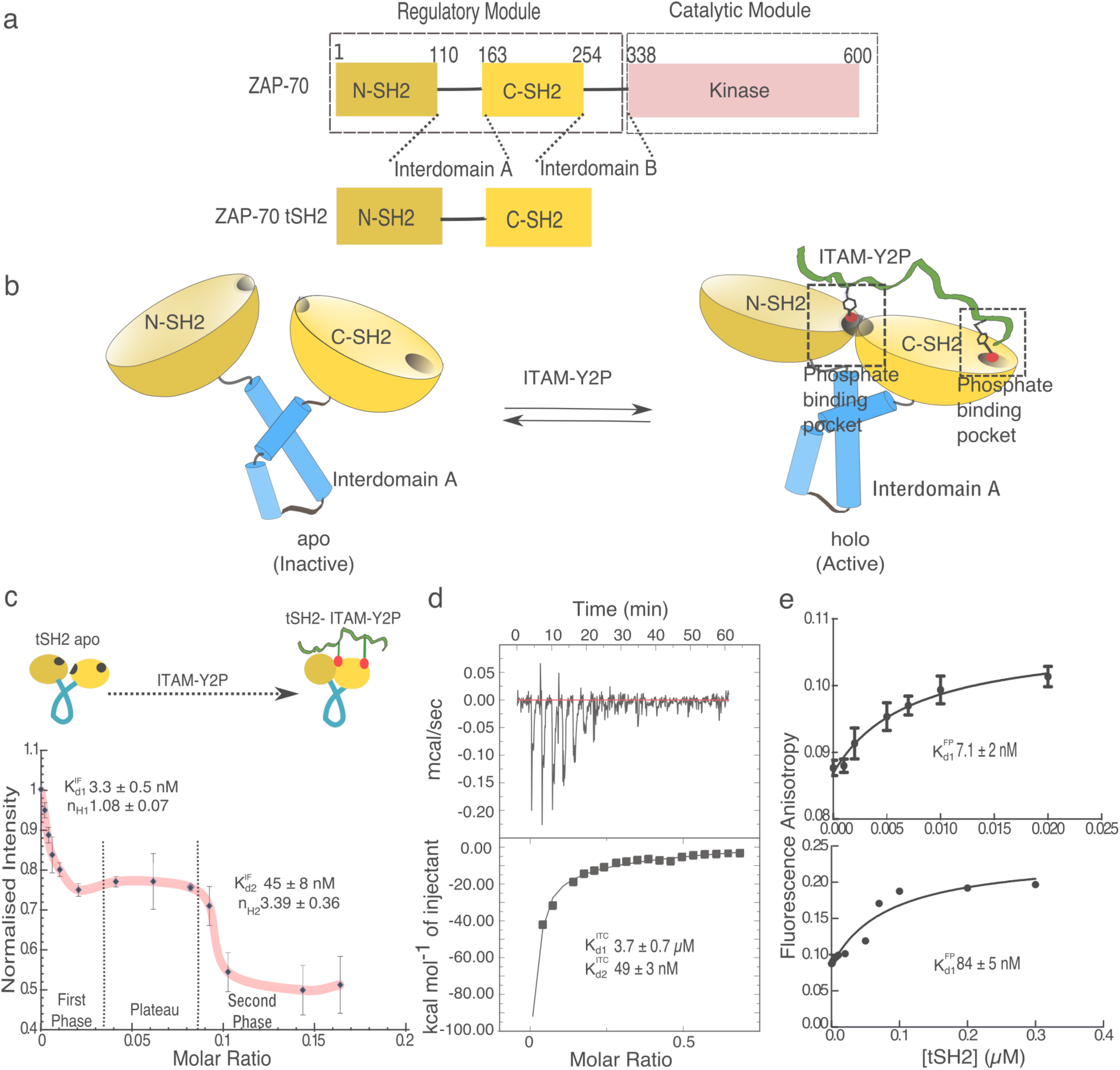
Binding of ZAP-70 tSH2 domain and doubly-phosphorylated ITAM-ζ1. (a) Schematic representation of domain architecture of full-length ZAP-70 and tSH2 domain used in this study. (b) Cartoon representation of tSH2-*apo* (unbound; PDB ID: 1M61) and tSH2-*holo* (doubly-phosphorylated ITAM bound; PDB ID: 2OQ1) structure of ZAP-70. The N-terminal SH2 domain (N-SH2), C-terminal SH2 domain, and respective phosphate-binding pocket are labeled. (c) Titration of doubly-phosphorylated ITAM-ζ1 peptide (ITAM-Y2P-ζ1) and tSH2 domain determined from the measurement of intrinsic tryptophan fluorescence at the indicated ligand to protein molar ratio. The solid red line is for guiding eyes. The dissociation constant for the first phase 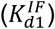 and the second phase 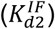 was determined from the curve-fitting to one-site specific binding model using Prism (Figure S1f and S1g). The Hill-coefficient (n_H_) was calculated from the Hill-plot (Figure S1c). (d) Representative isothermal titration calorimetry for the tSH2 domain and ITAM-Y2P-ζ1 peptide. (e) Binding of the Alexa Fluor 488-ITAM-Y2P-ζ1 peptide to the wildtype tSH2 domain was probed from the plot of fluorescence anisotropy *vs.* tSH2 domain concentration. Two independent experiments were performed at the indicated tSH2 domain concentration. The error bars indicate the standard deviation from three experiments.

The tSH2 domain of ZAP-70 binds with a high degree of selectivity and affinity to a conserved sequence of doubly-phosphorylated ITAM motif(30–34). The fundamental question of how does tSH2 domain, at the initial step, binds to the doubly-phosphorylated ITAM motif is not clearly known. Analysis of the crystal and NMR structures of an isolated tSH2 domain of ZAP-70 revealed that the phosphate-binding pocket of the C-SH2 domain is poised to bind first to the doubly-phosphorylated ITAM peptide(18). Through stochastic fluctuations, the two SH2 domains structurally reorient into geometrically close conformation forming the second phosphate-binding pocket (Figure 1b)(18). Alternatively, biochemical analysis and molecular dynamics simulation suggest that the N-SH2 domain may first bind to phosphotyrosine residue of ITAM peptide with low micromolar affinity followed by cooperative binding of second phosphotyrosine to the C-SH2 domain(11, 30, 32, 35).

Unlike tSH2 domain of spleen tyrosine kinase (Syk), a close homolog of ZAP-70 express in B-cells, unique aspect of ZAP-70 interaction to the TCR complex is the allosteric binding of the tSH2 domain to the doubly-phosphorylated ITAMs(11, 30, 36, 37) (Figure S1 and S2). The molecular mechanism of how the two SH2 domains of ZAP-70 allosterically cross-talk is not understood. A long-standing puzzle that is yet to be solved is how does a spontaneous mutation of W165C at the tSH2 domain reported in SKG mice alter the interaction of ZAP-70 to doubly-phosphorylated ITAM motifs at the membrane(17). W165, which is located far from the phosphate-binding pockets, impair the ZAP-70 activity causing defective thymic selection of developing T-cell leading to the development of chronic arthritis in the SKG mice.

In this paper, we investigated the interaction of doubly-phosphorylated ITAM-ζ1 (ITAM-Y2P-ζ1) peptide to the tSH2 domain of ZAP-70 and elucidated the mechanism of how the two SH2 domains are allosterically coupled. Our data showed a biphasic transition of the ZAP-70 tSH2 domain structure from an open to a closed state upon binding to doubly-phosphorylated ITAM-ζ1 peptide. Using molecular dynamics simulation, NMR spectroscopy, and biochemical analysis of different tSH2 domain mutants, we show that the C-SH2 domain binds first to the phosphotyrosine residue of the ITAM peptide. Following a plateau, the second phosphotyrosine residue of the ITAM peptide binds the N-SH2 phosphate-binding pocket. We deciphered an allosteric network, found only in ZAP-70, assembled by threading aromatic stacking interactions that connect N-SH2 and C-SH2 phosphate-binding pockets. The proposed model of allosteric network explained the molecular mechanism of altered interaction of W165C mutant of ZAP-70 and doubly-phosphorylated ITAM peptide in SKG mice.

## Results and discussion

### Fluorescence titration reveals biphasic binding of tSH2 domain of ZAP-70 to the doubly-phosphorylated ITAM peptide

The interaction of doubly-phosphorylated ITAM peptide and tSH2 domain of ZAP-70 has extensively studied, but the structural transition of tSH2 domain from the *apo* (ITAM-Y2P-ζ1 unbound state) to the *holo* state (ITAM-Y2P-ζ1 bound state) is poorly understood(6, 11, 18, 30–32, 34, 35) (Figure 1b). We used intrinsic tryptophan fluorescence (shown in Figure S1a and S1b) to monitor the structural transition of an isolated tSH2 domain of ZAP-70 upon binding to doubly-phosphorylated ITAM-ζ1 peptide. Titration of ITAM-Y2P-ζ1 to tSH2 domain quenches the tryptophan fluorescence producing a biphasic curve (Figure 1c and S1c) while transitioning from tSH2-*apo* to tSH2-*holo* state. The first phase showed a strong noncooperative binding [Hill-coefficient (n_H_) = 1.08 ± 0.07] with a dissociation constant 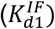 of 3.3±0.5 nM (Figure 1c, S1d and S1f), which is followed by second cooperative binding (n_H_ = 3.39 ± 0.36) of the doubly-phosphorylated ITAM-ζ1 peptide with 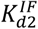 of 45 ± 8 nM (Figure S1d and S1g). The two binding events are interleaved by a plateau where no conformational changes were observed. We next tested the binding of ITAM-Y2P-ζ1 to the tSH2 domain of Syk by intrinsic tryptophan fluorescence spectroscopy (Figure 1a) (38, 39). As reported previously, the tSH2 domain of Syk binds uncooperatively (n_H_=1.09 ±0.01, 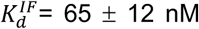) to the doubly-phosphorylated ITAM-ζ1 peptide, which undergoes a hyperbolic structural transition from *apo* to *holo* state(38) (Figure S2e).

To find out if the doubly-phosphorylated ITAMs follow a biphasic binding to the ZAP-70 tSH2 domain, we study the binding of doubly-phosphorylated ITAM-ζ1 peptide to tSH2 domain by fluorescence polarization and isothermal titration calorimetry (ITC) measurements. The titration of fluorescently labeled (Alexa-Fuore 488) ITAM-Y2P-ζ1 to the tSH2 domain also produces a biphasic curve with two dissociation constants of 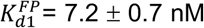 and 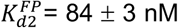 (Figure 1e and S1e). The ITC data that was fitted to two-site sequential binding model generating two dissociation constants (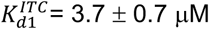 and 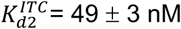) (Figure 1d).

In summary, we showed that the tSH2 domain of ZAP-70 binds to the doubly-phosphorylated ITAM-ζ1 peptide in a biphasic pattern with three distinct binding events correspond to strong, medium and weak dissociation constant regimes (Table 1). We observed that the first binding is strong low nano-molar (*K*_d_: 3-10nM) and uncooperative, second binding is weak micro-molar (*K*_d_: 2-4 μM) but positively cooperative to the third medium binding (*K*_d_: 50-80 nM). It is not clear how the tSH2 domain of ZAP-70, which has two phosphotyrosine binding pockets exhibit three binding events. A review of the literature shows that the measured dissociation constant matches with the published binding affinities for the tSH2 domain of ZAP-70 and doubly-phosphorylated ITAMs (Table S1). We noted that each of the binding affinity reported earlier could explain the biphasic binding when considered together. We next focused our effort to understand the mechanism of biphasic binding of doubly-phosphorylated ITAMs to the tSH2 domain. We begin by examining out of two phosphate-binding pockets which SH2 domain is binding first to the doubly-phosphorylated ITAM peptide (Figure 1b).

**Table 1:** Binding constant of tSH2 domain of ZAP-70 for ITAM-Y2P-ζ1

| Protein | Intrinsic Fluorescence<sup>a</sup> |  | Isothermal Titration Calorimetry<sup>b</sup> |  | Fluorescence Polarisation<sup>a</sup> |  |
| --- | --- | --- | --- | --- | --- | --- |
| | ($K_{d1}^{IF}$) | ($K_{d2}^{IF}$) | ($K_{d1}^{ITC}$) | ($K_{d2}^{ITC}$) | ($K_{d1}^{FP}$) | ($K_{d2}^{FP}$) |
| tSH2 wildtype | 3.3 ± 0.5 nM | 45 ± 8 nM** | 3.7 ± 0.7 μM | 49 ± 3 nM | 7.1 ± 2 nM | 84 ± 5 nM |
| F117A | 8.4 ± 1.8 nM | -- | 34.15 ± 8.79 μM | 483.35 ± 53.28 nM | 9 ± 2.1 nM | -- |
| W165C | 5.6 ± 1.6 nM | -- | 25.34 ± 0.99 μM | 939.44 ± 52.86 nM | 8 ± 1.9 nM | -- |
| R43P | 11 ± 2.5 nM | -- | -- | -- | 10 ± 3 nM | -- |

<sup>a</sup> Binding constant ( $K_d^{IF}$ , $K_d^{FP}$ ) for tSH2 wildtype and mutants are calculated using Prism from three independent experiments. <sup>b</sup> Binding constants ( $K_d^{ITC}$ ) are calculated using Origin from three independent experiments. \*\* The data was fitted to one-site specific binding with R<sup>2</sup> of 0.85, shown in figure S1f.

### Molecular dynamics simulation predicts the stronger binding of doubly-phosphorylated ITAM peptide to the C-SH2 domain

Our data suggest that the tSH2 domain of ZAP-70 has one strong and one weak phosphate-binding pocket. To examine out of two phosphate-binding pockets which one binds strongly or weakly to the phosphotyrosine residue of ITAM, we carried out molecular dynamics (MD) simulations of different ZAP-70 tSH2 domain structures (Figure 2a) and studied the time-dependent behavior. We begin by analyzing the average root-mean-square deviation (RMSD) that provides a qualitative measure of the protein structure and dynamics during the simulation. The average Cα RMSD value of the tSH2 domain bound to ITAM-Y2P-ζ1 (tSH2-*holo*) and ITAM-Y2P-ζ1 unbound (tSH2-*apo*) structures are 2.79 ± 0.28 Å and 6.44 ± 0.45Å, respectively (Figure 2b and S3a), suggests that the binding of doubly-phosphorylated ITAM peptide quenches the overall backbone dynamics of the tSH2 domain(21, 35). The structure in which either the N-SH2 (N-SH2^ITAM-YP^) or C-SH2 domain (C-SH2^ITAM-YP^) phosphate-binding pocket is occupied by phosphotyrosine residue of ITAM, deviates from the tSH2-*holo* structure with an average RMSD of 6.40 ± 0.44 Å and 5.45 ± 0.81Å, respectively. The N-SH2^ITAM-YP^ structure spontaneously adopts an open conformation and remains in the open conformation throughout the rest of the simulation trajectory. The C-SH2^ITAM-YP^ structure exhibits significant fluctuations of RMSD in the simulation trajectory (Figure 2b). We observed that the C-SH2^ITAM-YP^ structure undergoes a conformational transition between several states, including an open (at 12 ns and RMSD of 7.59 Å) and a close (at 40 ns and RMSD of 3.15 Å) conformation of the tSH2 domains, respectively (Figure 2e).

**Figure 2:**
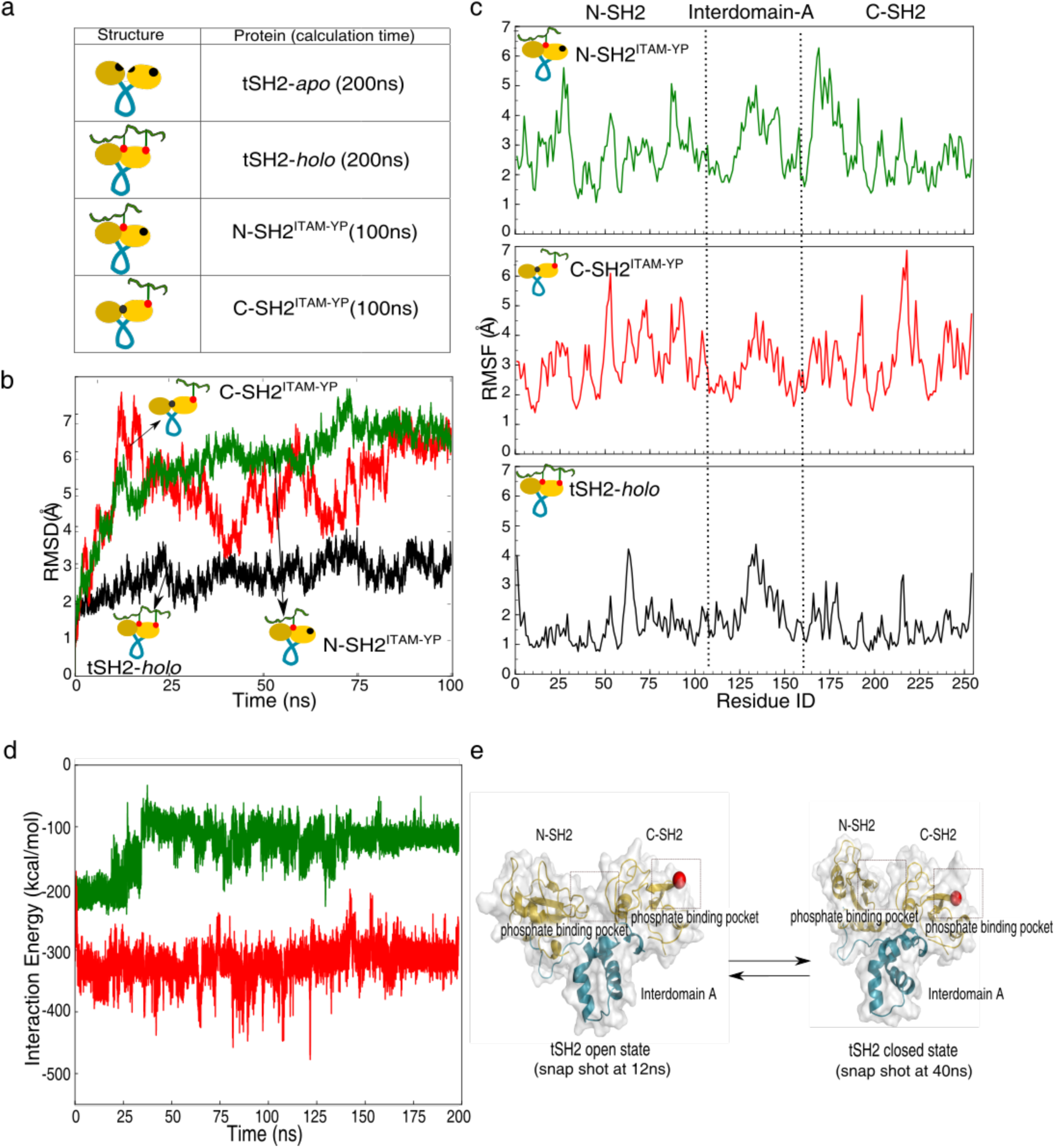
Structural evolution of the tSH2 domain of ZAP-70 and doubly-phosphorylated ITAM-ζ1 during MD simulation. (a) Schematic representation of different tSH2 domain constructs used in the MD simulation and their respective simulation time. (b) Cα root-mean-square deviation (RMSD) of the tSH2-*holo*, N-SH2^ITAM-YP^, and C-SH2^ITAM-YP^ structures are presented from the 100 ns simulation trajectory. (c) Average root-mean-square fluctuation (RMSF) for the 100 ns simulation in tSH2-*holo*, N-SH2^ITAM-YP^, and C-SH2^ITAM-YP^ structures are plotted against the residue number. (d) Interaction energy profile of the N-SH2 domain (green) and C-SH2 domain (red) with respective phosphotyrosine residue of ITAM-Y2P-ζ1 peptide in the tSH2-*holo* structure. (e) Representative snapshots of the tSH2 domain structures from the C-SH2^ITAM-YP^ trajectory corresponds to 12ns (high) and 40ns (low) time scale.

We analyzed the average root-mean-square fluctuations (RMSF) to elucidate the domain-specific dynamic behavior of the protein at the residue level. We observed an overall increase in RMSF for the tSH2-*apo*, N-SH2^ITAM-YP^, and C-SH2^ITAM-YP^ structures than the tSH2-*holo* state (Figure 2C). Map of residue-specific RMSF greater than 3.5 Å on the tSH2-*holo* structure (Figure S3c) revealed that the binding of phosphotyrosine residue to the N-SH2 phosphate-binding pocket leads to increase in structural flexibility of the C-SH2 phosphate-binding pocket. In the N-SH2^ITAM-YP^ simulation, the αA-helix at the C-SH2 phosphate-binding pocket is relatively more flexible compared to the tSH2-*holo* structure.

We next evaluated the theoretical binding affinity of the C-SH2 and the N-SH2 phosphate-binding pockets for the phosphotyrosine residue of ITAM in the tSH2-*holo* structure. The binding affinity was analyzed from the non-bonded interaction energy between the phosphotyrosine residue of ITAM and respective N-SH2 or C-SH2 domain, including the amino acid residues in the respective phosphate-binding pockets (Figure 2d). Our analysis indicates that the C-SH2 domain may bind phosphotyrosine residue with a stronger affinity than the N-SH2 domain. The average interaction energies are found to be −130.22 kcal/mol and −321.36 kcal/mol for the N-SH2 and C-SH2 phosphate-binding pockets, respectively. In summary, our simulation studies suggest that the C-SH2 domain may bind first with a strong affinity to the doubly-phosphorylated ITAM peptide leading to the formation of an encounter complex. We speculate that the encounter complex may enhance the structural dynamics of the tSH2 domain resulting in N-SH2 and C-SH2 domain to rearrange transiently in a closed conformation.

### C-terminal SH2 domain binds first to the doubly-phosphorylated ITAM peptide

To test our observations from the MD simulations, we studied the conformational rearrangement of the tSH2 domain of ZAP-70 upon binding to doubly-phosphorylated ITAM-ζ1 peptide by nuclear magnetic resonance (NMR) spectroscopy. The ^15^N-^1^H TROSY spectra of the tSH2-*holo* state produce well-dispersed peaks suggesting a folded and structurally homogenous protein (Figure S4). In the ^15^N-^1^H TROSY spectra of the tSH2-*apo* sample, we observed an overall decrease in the intensities and line broadening of several backbone amide peaks. The increased backbone dynamics for the tSH2-*apo* structure was reflected in the MD simulation and previously reported NMR studies of an isolated tSH2 domain(21, 35) (Figure S3). Thus, we considered a decrease in peak intensity due to line broadening as a hallmark for the tSH2-*apo* state.

To get insight into the sequential binding of doubly-phosphorylated ITAM peptide to the tSH2 domain, we titrate the ^15^N labeled tSH2-*apo* protein to a sample containing 1:2 mixture of ^15^N labeled tSH2: unlabeled ITAM-Y2P-ζ1 and recorded ^15^N-^1^H HSQC spectra (Figure 3a and S5). We observed that removal of doubly-phosphorylated ITAM decreases the overall intensity for the backbone resonances. The plot of residue number versus normalized intensities from each titration point suggests that the amino acid residues at N-SH2 phosphate-binding pocket line-broadens at a higher ligand to protein ratio (0.6) than C-SH2 domain (Figure 3a). Based on the plot of amino acid intensity versus the molar ratio of ITAM-Y2P-ζ1: protein (Figure S5), we classified the amino acid residues of the tSH2 domain into four groups (I to IV). The amino acid residue belongs to groups I, II, and III disappeared (due to line broadening) at a ligand to protein ratio of 0.6, 0.4, and 0.2, respectively. All other residues that did not line-broaden at ligand to protein ratio of 0.2 were considered into group IV. Analysis of the backbone amide intensity for the amino acid residues at the N-SH2 and C-SH2 phosphate-binding pockets show a clear distinction (Figure 3b and d). The amino acid residues, R19, R39, C41, L42, R43, S44, and H60 that are in close contact with the phosphotyrosine residue of ITAM at the N-SH2 phosphate-binding pocket are clustered into group I and II (Figure 3b and d) with micromolar dissociation constants (Figure S5). The disappearance of backbone resonances for the amino acid residues at the N-SH2 phosphate-binding pocket at a relatively higher concentration of doubly-phosphorylated ITAM-ζ1 peptide indicates that the N-SH2 phosphate-binding pocket represents the weaker phosphate-binding site. Whereas, the amino acid residue at the C-SH2 phosphate-binding pocket, T171, L190, R192, R194, S203, Y213, and H212 were clustered into group III and IV representing the strong affinity site. Residue R194 and R192 that interacts with the phosphotyrosine residue of ITAM also showed ligand depended on chemical shift changes (Figure 3c).

**Figure 3:**
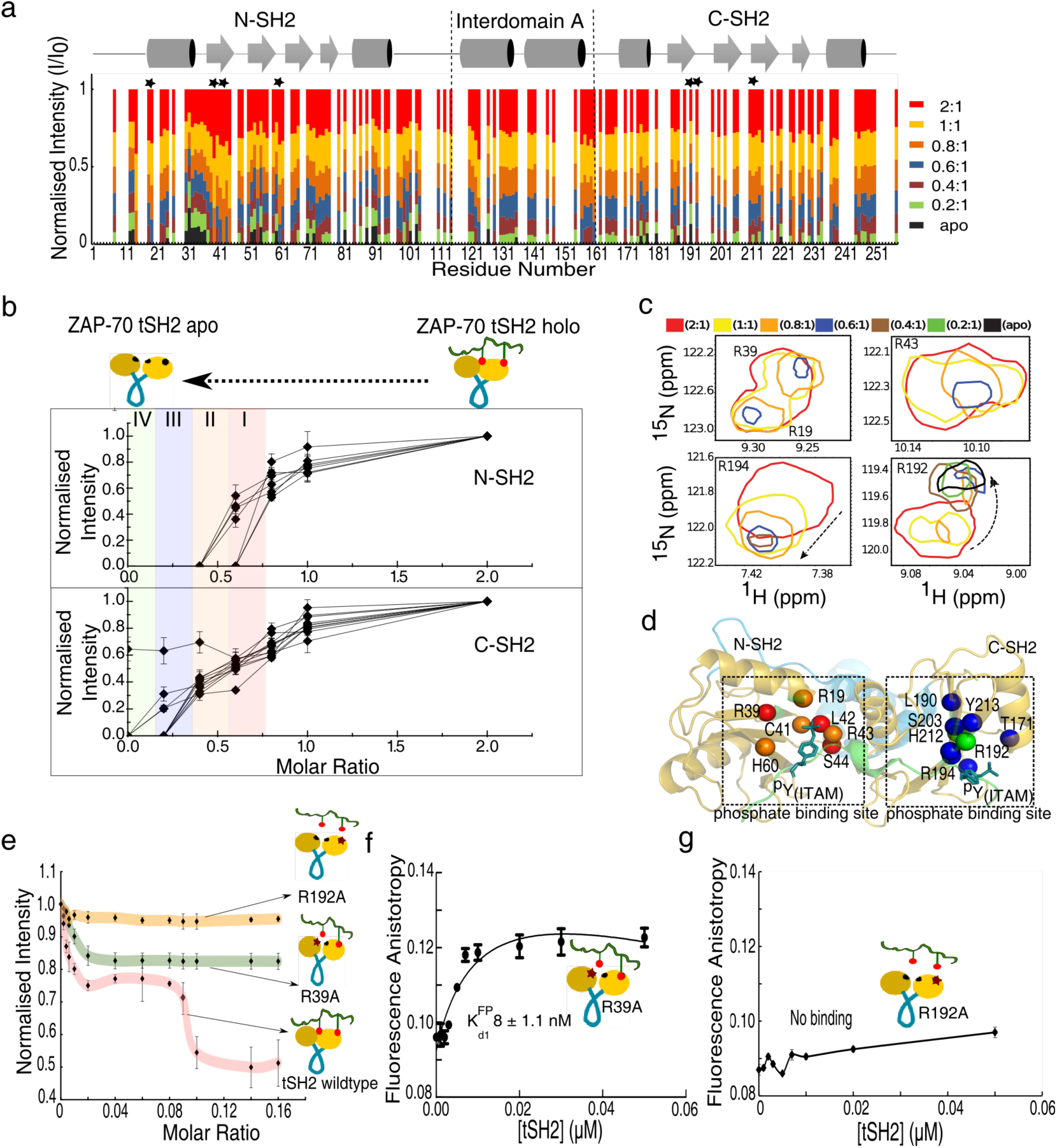
Titration of ZAP-70 tSH2 domain and doubly-phosphorylated ITAM-ζ1 by NMR spectroscopy. (a) Normalized intensity of the backbone amide resonances measured from each NMR titration experiment (color-coded) is plotted as a function of residue number. The intensity was normalized by the intensity of the respective amino acid residues measured with a sample made up of 2:1 ligand to protein ratio. The secondary structure of the tSH2 domain in the *holo* state is shown at the top. Amino acid residues at the phosphate-binding pocket interacting with the ITAM-Y2P-ζ1 phosphate group are indicated by a star. (b) The normalized intensity of the backbone amide region for the amino acid residues at the N-SH2 and C-SH2 phosphate-binding pockets are plotted against the ligand to protein molar ratio. At the top panel, amino acid residues R19, R39, C41, L42, R43, S44, and H60 from the N-SH2 phosphate-binding pocket are plotted. In the bottom panel amino acid residues T171, L190, R192, R194, S203, H212, and Y213 at the C-SH2 phosphate-binding pocket are shown. The vertical color represents the four class of amino acid residues described in the text and figure S5. (c) Representative cross-section showing the overlapped ^15^N-^1^H HSQC spectra from the NMR titration experiments (color-coded). The direction of chemical shift change is shown by the arrow. (d) The amino acid residues (shown as a sphere) from panel b is mapped on the tSH2-*holo* structure (PDB ID: 2OQ1). The color code represents the class of each amino acid, as described in panel b. (e) Binding of ITAM-Y2P-ζ1 to the R39A and R192A mutant of the tSH2 domain of ZAP-70 determined from the intrinsic tryptophan fluorescence is plotted against the ligand to protein molar ratio. The solid-colored line is for guiding eyes. (f) and (g) is the plot of fluorescence anisotropy of the Alexa Fluor 488-ITAM-Y2P-ζ1 peptide against the concentration of R39A and R192A mutated tSH2 domain, respectively.

We tested the ability of the domain-specific mutants (R39A and R192A at the N-SH2 and C-SH2 phosphate-binding pockets, respectively) of the tSH2 domain to bind doubly-phosphorylated ITAM-ζ1 peptide by fluorescence spectroscopy (Figure 3e and S6). Indeed, mutation of R192A at the C-SH2 phosphate-binding pocket impaired the ITAM-Y2P-ζ1 peptide binding to the tSH2 domain and did not show the biphasic structural transition from *apo* to *holo* state (Figure 3e and g). Whereas, the R39A mutant binds uncooperatively (n_H_ = 1.3 ± 0.12) to ITAM-Y2P-ζ1 with low-nanomolar binding affinity (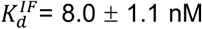 and 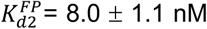) (Figure 3e, and S7). Our data indicate that the binding of phosphotyrosine to the C-SH2 phosphate-binding pocket is imperative for the subsequent phosphotyrosine binding to the N-SH2 phosphate-binding pocket.

To test if the binding of doubly-phosphorylated ITAM peptide to the C-SH2 domain alone could induce close conformation, we evaluated the conformation of the tSH2 domain mutants by measuring acrylamide quenching of tryptophan fluorescence in the *apo* and the *holo* states (Figure S6)(40). The wildtype tSH2-*apo* state showed concentration-depended acrylamide quenching of tryptophan fluorescence yielding a slope (Stern-Volmer quenching constant, *K*_*sv*_) of 0.044 ± 0.003 μM^−1^, suggesting open conformation of tSH2 domain is amenable for acrylamide quenching. The closed conformation of the tSH2-*holo* state shields the tryptophan from acrylamide quenching (decreased the *K*_*sv*_ to 0.015 ± 0.001 μM^−1^). At the plateau of the tryptophan fluorescence titration curve (Figure 1c), the tSH2 structure exhibits an intermediate *K*_*sv*_ value of 0.027 ± 0.002 μM^−1^, indicating that the tSH2 domain may exist in a dynamic equilibrium between a closed and an open state (Figure S6b). The R192A mutant of the tSH2 domain that does not bind to the ITAM-Y2P-ζ1 peptide remains in an open conformation (*K*_*sv*_ = 0.046 ± 0.001 μM^−1^) even in the presence of the peptide (Figure S6 and 2b). Whereas the intermediate *K*_*sv*_ value (0.025 ± 0.003 μM^−1^) for the tSH2-*holo*^R39A^ sample indicates that the tSH2 domain could adopt closed conformation transiently when the C-SH2 phosphate-binding pocket is bound to the phosphotyrosine residue of ITAM (Figure S6 and 2b). Together our NMR data, MD simulation, and biochemical analysis of tSH2 domain mutants suggest that the binding of the phosphotyrosine to the C-SH2 domain transiently aligns the two SH2 domains into closed proximity in a geometrical arrangement facilitating the formation of second phosphate-binding pocket (Figure 2). However, these data do not explain how the two SH2 domains cross-talk during binding to the doubly-phosphorylated ITAM peptide.

### Aromatic-aromatic interaction constitute an allosteric hot-spot that connects N- and C-SH2 domains

To understand the structural coupling between the C- and N-SH2 domain during doubly-phosphorylated ITAM-ζ1 binding, we analyzed the backbone amide chemical shift differences for the tSH2 domain observed at each titration point. Figure 4 shows the plot of residue numbers versus the difference in the compounded chemical shifts (ΔΔCCS) observed. The significant ΔΔCCS observed in each titration step was mapped onto the tSH2-*holo* structure (Figure 4B). In general, we observed coupled chemical shift changes at both the phosphate-binding sites and at the interdomain-A region. The ΔΔCCS due to structural rearrangement at the N-SH2 domain propagates into the phosphate-binding pocket at the C-SH2 domain and to the interdomain-A. For example, at a ligand to protein ratio of 0.8 and 0.6, we observed chemical shift perturbation for key amino acid residues at the N-SH2 phosphate-binding pocket, which induces chemical shift change for the T171, R192, and R194 at the C-SH2 domain phosphate-binding pocket (Figure 4a and b). Thus, our NMR data suggest that the binding of phosphotyrosine of ITAM at the N-SH2 domain remodels the structure of the C-SH2 phosphate-binding pocket.

**Figure 4:**
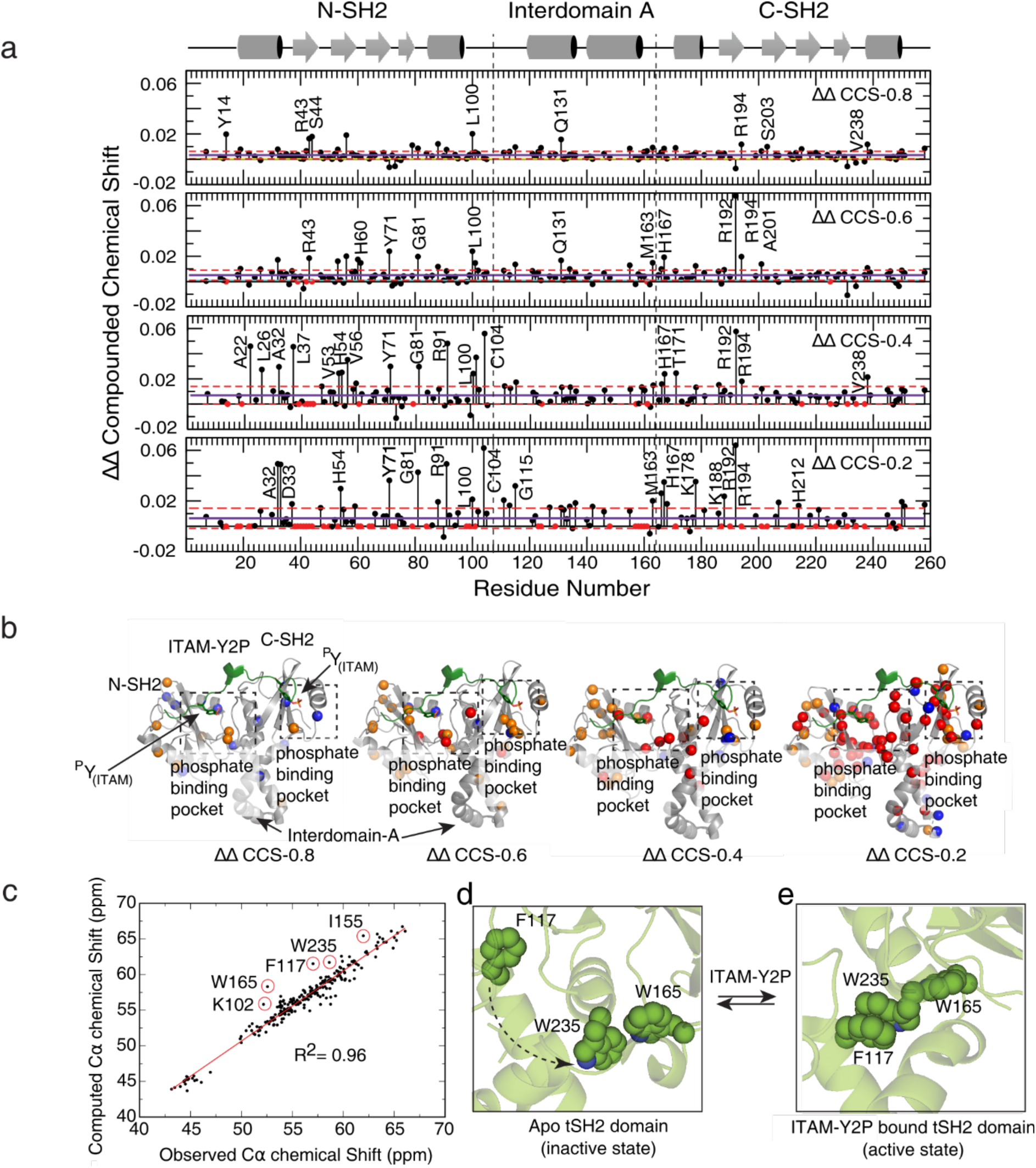
NMR chemical shift analysis of the tSH2 domain of ZAP-70 upon binding to doubly-phosphorylated ITAM-ζ1. (a) ΔΔcompounded chemical shift (ΔΔCCS) change of backbone amid resonances of tSH2 domain of ZAP-70 observed during the titration of ITAM-Y2P-ζ1 peptide is plotted against the residue number. Each panel represents the chemical shift change observed at the indicated ligand to protein ratio. The solid horizontal line and the broken red line represent the average chemical shift change and the standard deviation, respectively. Residues showing ΔΔCCS more than the average + Std are labeled in each panel. The residues disappear due to line-broadening during each titration step are shown as a red circle. At the top, the secondary structure of the tSH2-*holo* state is shown. The vertical dashed line represents the domain boundaries. (b) Residues experiencing chemical shift changes or line broadening are mapped on to the crystal structure of the tSH2 domain bound to ITAM-Y2P-ζ1 (PDB ID: 2OQ1). The orange and blue sphere represents amino acid residues that showΔΔCCS more than average + Std and within the Std, respectively. Residues that line-broadens beyond detection are colored red. (c) Correlation plot of Cα chemical shift of tSH2 domain bound to ITAM-Y2P-ζ1 peptide measured from the NMR experiments and calculated from the crystal structure of the tSH2-*holo* state. (d) and (e) is the conformation of the aromatic residues at the proposed allosteric hot-spot in the tSH2-*apo* (PDB ID: 1M61) and the tSH2-*holo* (PDB ID: 2OQ1) structures, respectively.

To explain the chemical shift changes observed for residues at the interface of the interdomain -A and C-SH2 domain, we evaluate the backbone structure of the tSH2-*holo* state from crystallography and NMR spectroscopy (Figure 4c). We compare the Cα chemical shifts of the tSH2-*holo* state measured from the NMR experiments to the Cα chemical shifts predicted from the crystal structure (PDB: 2OQ1). Cα chemical shift is influenced by the backbone conformation of amino acids in a protein, thus provide an excellent parameter to compare the two structures(41). We observed an overall agreement between (R^2^=0.96) the Cα chemical shift from the crystal structure and NMR experiments (Figure 4c). However, few residues located at the interface of the N-SH2 domain, interdomain-A, and C-SH2 domain, namely F117, W165, and W235 stand out as an outlier. In the *holo* state, F117, W165, and W235 are locked in close conformation stabilized by aromatic-aromatic interaction (Figure 4e). Release of doubly-phosphorylated ITAM peptide from the tSH2 domain breaks the F117-W235 interaction and reorients the W235 and W165 to adopt an open conformation. Based on the NMR data and analysis of the crystal structures we hypothesized that the F117, W165, and W235 constitute an allosteric hot-spot that couples the two SH2 domains through a network of noncovalent interaction during ligand binding.

### A network of noncovalent interaction couples C- and N-terminal SH2 domains through the allosteric hot-spot

To elucidate a network of noncovalent interaction connecting the C-SH2 and the N-SH2 domains through allosteric hot-spot residues, we performed a comparative analysis of the residue interaction network (RIN) of tSH2-*apo* (PDB ID: 1M61) and tSH2-*holo* (PDB ID: 2OQ1) structure of ZAP-70 (Figure 5)(42, 43). The residue interaction network map was constructed by using the Rin-analyzer plugin(42, 44) of Cytoscape(45, 46). Table S4 summarizes the network parameters for the *apo* and *holo* tSH2 domain. The residue interaction network for the tSH2-*apo* is comprised of 259 nodes that are connected by 2630 noncovalent interaction (NCI) edges. Binding of doubly-phosphorylated ITAM remodels the residue interaction network in the tSH2-*holo* structure, where 273 nodes are now connected by 3305 noncovalent interaction edges. A plot of residue number versus node degree shows an overall increase in neighborhood connectivity for the tSH2-*holo* structure (Figure 5A).

**Figure 5:**
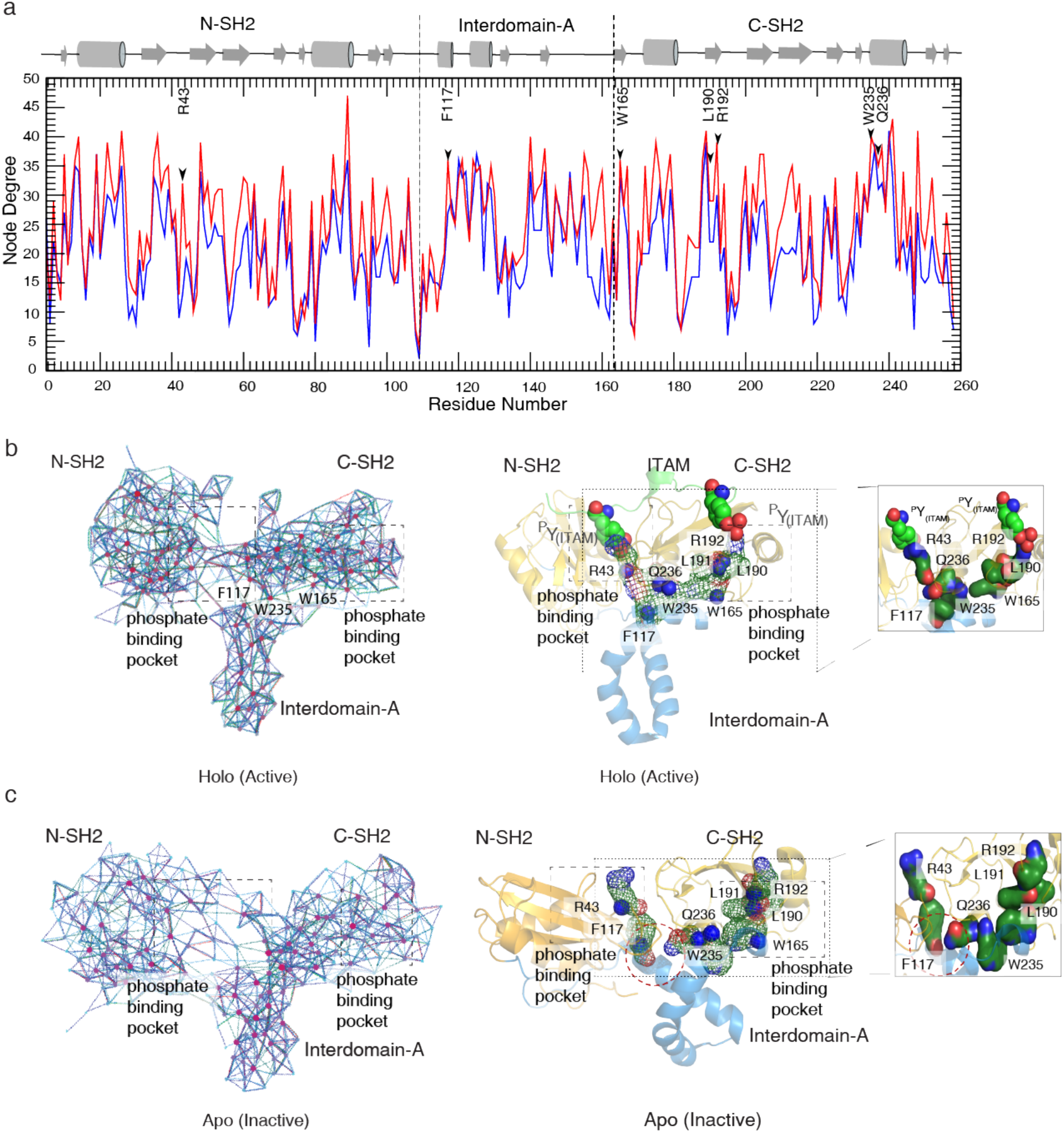
Non-covalent residue interaction networking in the tSH2 domain of ZAP-70. (a) The node degree for the tSH2 domain in the *apo* (blue line) and *holo* (red) conformation is plotted against the residue number. The secondary structure of the tSH2-*holo* state is shown at the top. Vertical broken lines indicate the domain boundaries. (b) and (c) are the schematic representation of the residue interaction network of the tSH2-*holo* (PDB ID: 2OQ1) and tSH2-*apo* structure (PDB ID: 1M61), respectively, visualized in Cytoscape. Each amino acid in the structure is represented as a node, and the non-covalent interaction connecting two nodes is represented as lines (edges). The amino acid residues with high node degree (hub residues) are highlighted as red circles. On the right side of each panel, the shortest residue interaction network connecting the two SH2 domains are mapped on the crystal structure of tSH2-*holo*, and tSH2-*apo* states, respectively.

We began our analysis with the tSH2-*holo* structure and searched for the shortest residue interaction pathway involving a minimum number of steps (amino acids) connecting the two-phosphate binding pockets through the allosteric hot-spot residues. As shown in figure 5b, the network initiates with R192 at the phosphate-binding pocket of the C-SH2 domain, which is connected to the W235 and W165. W235, in turn, is connected to F117 by π-π aromatic stacking interaction that finally converged to R43 at the N-SH2 phosphate-binding pocket. In the *holo*-state W235 has the highest node degrees is sandwiched between F117 and W165. Which suggests that W235 might function as an allosteric switch (nodal hub) that couples the two SH2 domains during ITAM binding. In the *apo-*state, the F117-W235 π-π aromatic stacking interaction is broken, which might uncouple the allosteric network between C-SH2 and N-SH2 domains (Figure 5b).

### Mutation in the allosteric network uncouples doubly-phosphorylated ITAM binding to C-SH2 domain from N-SH2 domains

To evaluate the importance of the allosteric-network in the tSH2 domain of ZAP-70 during doubly-phosphorylated ITAM binding, we studied the overall structure and dynamics of four in-silico mutants tSH2-*holo*^W165C^, tSH2-*holo*^F117A^, tSH2-*holo*^R43A^ and tSH2-*holo*^R43P^ (Figure 6a). All the tSH2 domain in-silico mutants used in the molecular dynamics simulations were prepared on the tSH2-*holo* structure (PDB ID: 2OQ1). Analysis of the structures from the molecular dynamics trajectory shows an overall increase in RMSD of 2.79 Å, 5.12 Å, 4.84 Å, 6.85 Å, 4.73 Å for tSH2-*holo*, tSH2-*holo*^W165C^, tSH2-*holo*^F117A^, tSH2-*holo*^R43A^ and tSH2-*holo*^R43P^, respectively (Figure S8). The higher RMSD for the mutants suggest that the conformation of the mutated tSH2 domain structures deviate from the wildtype tSH2-*holo* structure. We also observed an overall increase in structural flexibility (RMSF) in the N-SH2 and C-SH2 domains of the mutated tSH2-*holo* structure in comparison to the wildtype (Figure 6b).

**Figure 6:**
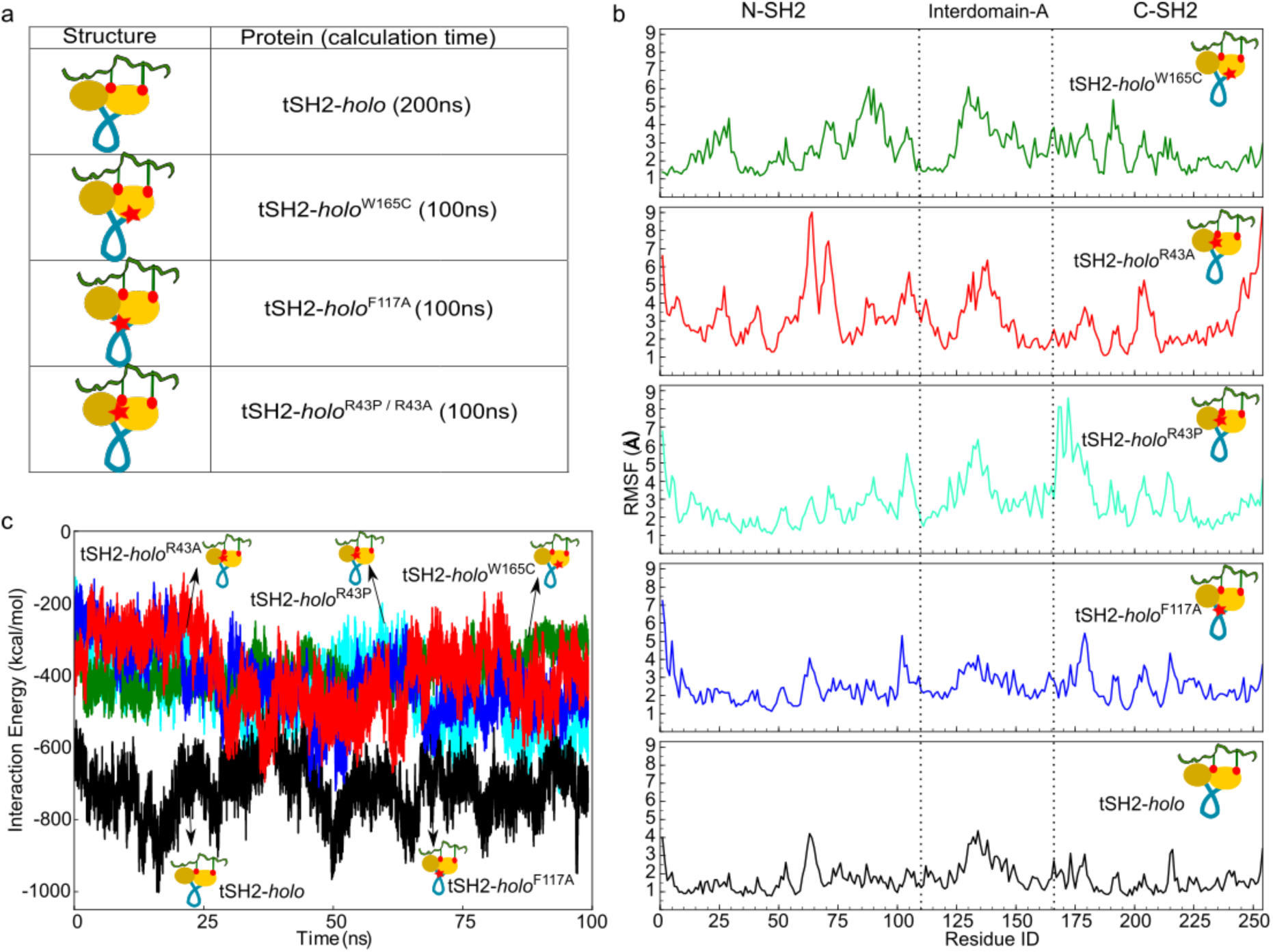
Structural evaluation of allosteric-network in the tSH2 domain of ZAP-70 by MD simulation. (a) Schematic representation of different systems considered for simulation and their respective simulation time scales. (b) Average root-mean-square fluctuation (RMSF) for the 100 ns simulation in tSH2-*holo* and mutated tSH2 structures bound to the ITAM-Y2P-ζ1 peptide are plotted against the residue number. (c) Interaction energy profile between the ITAM-Y2P-ζ1 peptide and tSH2-*holo*,tSH2-*holo*^W165C^, tSH2-*holo*^F117A^, tSH2-*holo*^R43A^ and tSH2-*holo*^R43P^ mutant structures are plotted against the simulation time, respectively.

To determine if the residue interaction network [R43-Q236-F117 -W235-W165-L191 (L190)-R192] coupling the two SH2 domains is present throughout the simulation trajectories, we measured the time-dependent pairwise distance between the residues with the representative side-chain atoms (Figure S9). We observed that the network connectivity was maintained throughout the simulation trajectory for the wildtype tSH2-*holo* structure (Table S3 and Figure S9a). However, we noted that during the simulation, Q236 rearranges in a stacking position between F117 and W235, providing stability to the network. In the network-mutants, tSH2-*holo*^W165C^, tSH2-*holo*^F117A^, tSH2-*holo*^R43A^ and tSH2-*holo*^R43P^, residue interaction-network connecting the two SH2 domains was significantly destabilized and broken (Figure S9b-e). The mutation increases the average pairwise distance between the key amino acid residues in the network (Table S3). The increase in structural flexibility along with destabilization of residue interaction-network by the mutants (W165C, F117A, R43A, and R43P) indicates that the network-mutation might alter the binding of the doubly-phosphorylated ITAMs to the tSH2 domain.

We next investigate the strength of wildtype tSH2-*holo* and the mutated tSH2 structures to bind the doubly-phosphorylated ITAMs from the simulation trajectories of the respective system (Table S2). The average interaction energies were found to be −713.09 kcal/mol, −390.54 kcal/mol, −418.52 kcal/mol, −396.51 kcal/mol, −448.67 kcal/mol for tSH2-*holo*, tSH2-*holo*^W165C^, tSH2-*holo*^F117A^, tSH2-*holo*^R43A^ and tSH2-*holo*^R43P^ respectively (Figure 6c). Our analysis of MD trajectory suggests that mutation of the allosteric network residue may impair the binding of doubly-phosphorylated ITAM peptide to the tSH2 domain.

To test the role of the proposed allosteric network, we made three mutation R43P, F117A, and W165C in the tSH2 domain background. We probed the affinity of the network mutant to bind doubly-phosphorylated ITAM-ζ1 peptide and the structural transition of the tSH2 domain to close state, by fluorescence spectroscopy and ITC experiments. The titration of doubly-phosphorylated ITAM-ζ1 peptide followed by intrinsic tryptophan fluorescence of the tSH2 domain shows that the R43P and W165C mutation does not induce the biphasic structural transition of tSH2 domain upon binding to the peptide (Figure 7a). For the F117A mutant, doubly-phosphorylated ITAM-ζ1 binding does quench the tryptophan fluorescence, but the biphasicity was significantly altered. We noted that for all the three network mutants the strong uncooperative binding of the doubly-phosphorylated ITAM-ζ1 to the C-SH2 domain was preserved (Figure 7 and S10c, Table 1). We could not determine any medium or weak binding of doubly-phosphorylated ITAM-ζ1 peptide to the mutated tSH2 domain by fluorescence polarization measurements. However, we observed in the ITC experiments the medium, and the weak binding of phosphotyrosine residue of ITAM motifs are significantly altered for F117A and W165C mutants (Table 1 and Figure 7c). We could not detect any binding of doubly-phosphorylated ITAM-ζ1 to the R43P mutant by ITC (Figure 7c). The acrylamide quenching for the tSH2-*holo*^R39P^ and the tSH2-*holo*^W165C^ shows that (Figure S10) these two mutants could not adopt the closed conformation. We observed significant shielding (*K*_*sv*_ = 0.019 μM^−1^) for the tSH2-*holo*^F117A^, indicating that the F117A mutation still allows the formation of the closed conformation, but the mutant significantly impaired allosteric coupling between the two SH2 domains. Together the MD simulation and biochemical evaluation of tSH2 domain mutants suggest that the mutation of the allosteric hot-spot residue does not perturb formation of the encounter complex between the C-SH2 domain and the phosphotyrosine residue of ITAM motif. The allosteric network mutant uncouples subsequent binding of the doubly-phosphorylated ITAM to the N-SH2 phosphate-binding pocket. The ITC data suggests that the allosteric mutants impose a thermodynamic penalty on the tSH2 domain to adopt a closed conformation upon doubly-phosphorylated ITAM binding (Table S5).

**Figure 7:**
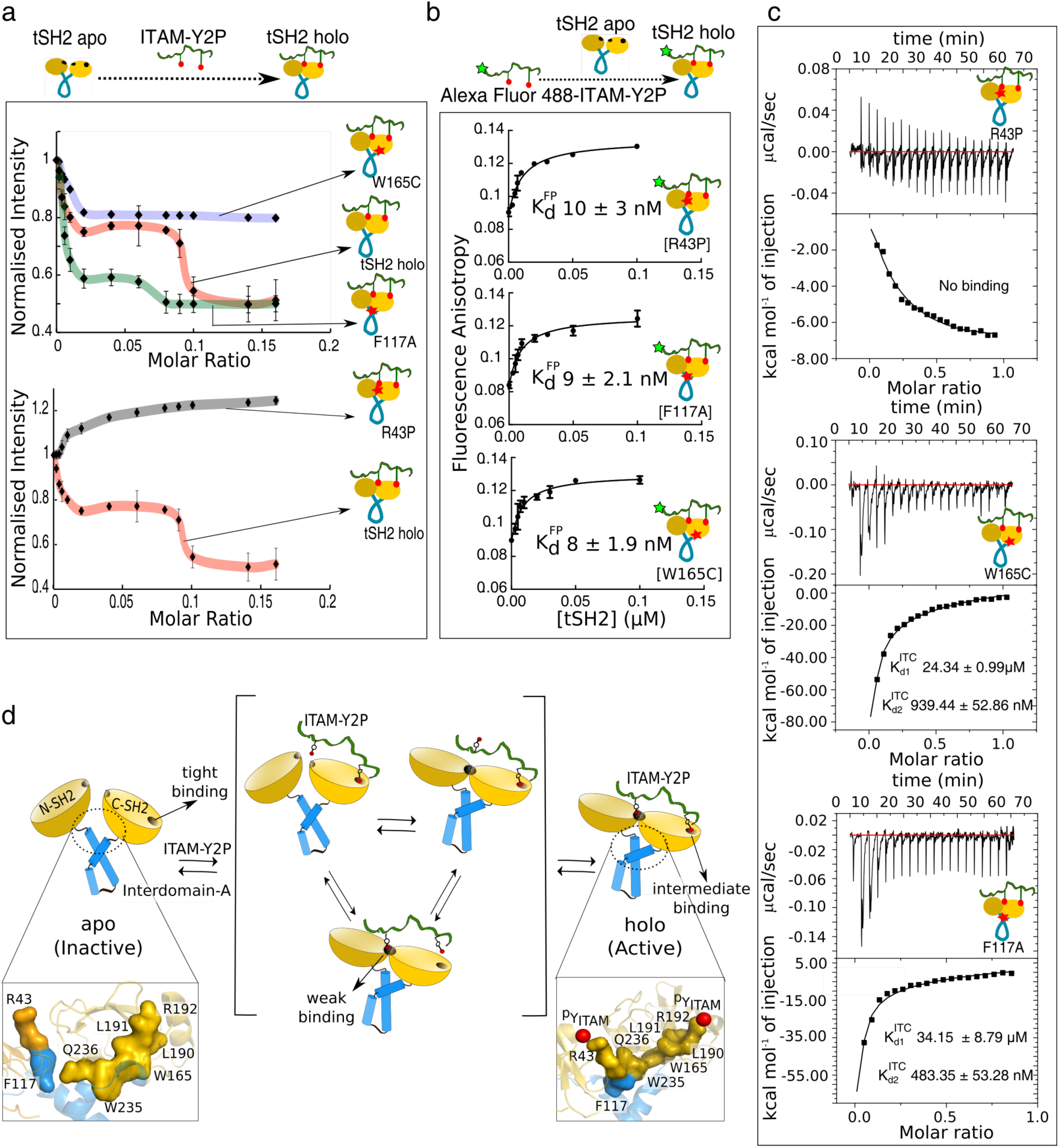
Effect of the allosteric-network mutant of ZAP-70 tSH2 domain on the binding of doubly-phosphorylated ITAM. (a) Comparative binding analysis of ITAM-Y2P-ζ1 and tSH2 domain mutants determined from the change in intrinsic tryptophan-fluorescence of the tSH2 domain during titration of ITAM-Y2P-ζ1 peptide. The normalized intensity for the W165C and F117A (top panel) and R43P (bottom panel) mutant of the tSH2 domain is plotted against the molar ratio of ligand to the protein. The solid-colored line is for guiding the eyes. The dissociation constant 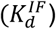 and the Hill-coefficient (n_H_) was obtained by fitting the data to one-site binding model implemented in Prism (Figure S10c) (b) Binding of Alexa Fluor 488-ITAM-Y2P-ζ1 peptide to R43P (top panel), F117A (middle panel) and W165C (bottom panel) mutant of tSH2 domain was probed from the plot of fluorescence anisotropy as a function of increasing tSH2 domain concentration. (c) Representative isothermal titration calorimetry profile for the titration of ITAM-Y2P-ζ1 peptide to R43P, W165C, and F117A mutant of tSH2 domain respectively. (d) Schematic representation of the model for the binding of ITAM-Y2P-ζ1 to the tSH2 domain of ZAP-70. In the inset, allosteric network residues in the tSH2-apo and tSH2-holo states are shown. The binding affinity of the N-SH2 and C-SH2 domains for the ITAM-Y2P-ζ1 are indicated.

We evaluated the stability of each of the structures (*holo* and *apo* conformation) of wildtype and mutated tSH2 domain by CD spectroscopy (Figure S11). As expected, *apo*- and the *holo-* state of the wildtype tSH2 domain represents the lowest and highest thermally stable conformation(32), respectively. All the tSH2 domain mutants in the *apo-*state clustered together with wildtype tSH2-*apo* structure, indicating the mutation did not change the overall stability of the proteins. The tSH2-*holo*^R192A^ mutant, which did not bind to doubly-phosphorylated ITAM-ζ1 peptide, exhibit similar thermal stability as the tSH2-apo. The tSH2-*holo*^R39A^ mutant that has a functional C-SH2 phosphate-binding pocket showed intermediate stability (Figure S11). The thermal stability of the *holo*-state for the wildtype and the allosteric network mutants of the tSH2 domain agrees with the simulation data. We noted that all the three allosteric mutants tSH2-*holo*^R43P^, tSH2-*holo*^W165C^, and tSH2-*holo*^F117A^ showed intermediate stability and clustered along with tSH2-*holo*^R39A^ mutant.

## Conclusion

Although Syk and ZAP-70 share high sequence similarity, similar domain architecture and activated by the conceptually same mechanism(11, 19–21, 37, 39, 47, 48), yet these two kinases recruit to the membrane by a fundamentally different mechanism. In contrast to Syk, the tSH2 domain of ZAP-70 undergoes a biphasic structural transition while binding to the doubly-phosphorylated ITAM peptide. In the first phase, phosphotyrosine residue of ITAM binds to the C-SH2 phosphate-binding pocket of the tSH2 domain with a low-nano molar affinity (*K*_d_: 3-10 nM) leading to the formation of an encounter complex (Figure 7d, 2d and 3d-g). The encounter in turn complex structurally couples the binding of second phosphotyrosine residue of ITAM peptide to the N-SH2 phosphate-binding pocket by transiently adopting a closed conformation of tSH2 domain (Figure 2b, 2e, S6, and 7d). The NMR chemical shift analysis and MD simulation data indicates that the second phosphotyrosine binding to the N-SH2 phosphate-binding pocket remodels the structure and dynamics of the C-SH2 phosphate-binding pocket, possibly to a medium (*K*_d_: 50-80 nM) affinity site (Figure 2c, S3c, and 4a). Therefore, at lower concertation of doubly-phosphorylated ITAM peptide, the second phosphotyrosine binding to the N-SH2 domain may release the phosphotyrosine residue from the C-SH2 phosphate-binding pocket, resulting in a plateau during the intrinsic tryptophan fluorescence experiment (Figure 1c and b). To adopt a stable tSH2-*holo* structure requires a reorientation of the aromatic residues F117, W165, and W235 into a stacking interaction, which imposes a higher energetic penalty (Figure S11 and Table S5). Finally, when the doubly-phosphorylated ITAM concentration builds up (to 50-80 nM), the C-SH2 domain spontaneously binds to the phosphotyrosine residue. The allosteric binding of the doubly phosphorylated ITAM peptide to the N-SH2 and C-SH2 phosphate-binding pockets are coupled through the residue interaction network that stabilizes the closed conformation of the tSH2-*holo* structure (Figure 7d).

Our proposed model of allosteric coupling in the tSH2 domain of ZAP-70 explains the altered binding of ZAP-70^W165C^ to doubly-phosphorylated ITAM found in SKG mice(17), which underlines the biological significance of the proposed allosteric network. The W165 residue is central to the residue interaction network that couples the (Figure 4 d-e and 7d) C-SH2 and N-SH2 phosphate-binding pockets through the F117-W235 interaction. Mutation of W165C breaks the residue interaction network and decouples the encounter complex from the subsequent phosphotyrosine binding (Table 1). We observed that tSH2-*holo*^W165C^ could not adopt a closed conformation like wildtype tSH2-*holo* structure because the mutation imposes a higher thermodynamic penalty on the tSH2 domain to adopt a closed conformation upon binding to doubly-phosphorylated ITAM (Table S5) (Figure S10b). Unlike Y126 in the interdomain-A that negatively regulates ITAM binding is conserved in both Syk and ZAP-70(28, 49–52). The proposed allosteric mechanism is a hallmark of ZAP-70 signaling that may provide an added regulatory mechanism essential for T-cell development and proliferation.

## Materials and Methods

Details of materials and method comprising of the Fluorescence experiments, Isothermal Titration Calorimetry, NMR spectroscopy, MD simulation can be found in Supplementary file.

## Supporting information

Gangopadhyay_MannaEtal_SI

## Author Contributions

The manuscript was written through contributions of all authors. All authors have given approval to the final version of the manuscript. RD, KG and SK performed the NMR experiment and data analysis. KG, SR and SC carried out the fluorescence spectroscopic and biochemical studies. OD performed the RIN analysis. AG and BM carried out the MD simulations. RD, AG, KG and BM wrote the manuscript.

## Acknowledgements

Authors are thankful to Prof. John Kuriyan and Prof. David E. Wemmer for access to the 900 MHz NMR spectrometer at the University of California, Berkeley. Dr. Jeffrey G. Pelton and Dr. Patrick R. Visperas at the University of California, Berkeley for NMR data collection and sample preparation. Authors thank Prof. Gautam Basu and Mr. Barun Majumder at Bose Institute, India for access to 700 MHz NMR spectrometer. Authors are thankful to Dr. Ashima Bhattacharjee, Dr. Pradip K. Tarafdar, and Prof. Pradipta Purkayastha for access to ITC and fluorimeter. Authors thank Prof. Giuseppe Melacini and Prof. Maitrayee DasGupta for helpful discussion. The authors thanks research funding from IISER Kolkata, infrastructural facilities supported by IISER Kolkata and DST-FIST (SR/FST/LS-II/2017/93(c)). This work is supported by grant from SERB (ECR/2015/000142) and DBT Ramalingaswami Fellowship (BT/RFF/Re-entry/14/2014) to RD. This work was supported by a research grant from the DST (No. ECR/2016/001096) and DBT-Ramalingaswami Re-entry fellowship (No.BT/RLF/Re-entry/06/2013) to A.G.

