## Supplementary material for "An allosteric hot spot in the tandem-SH2 domain of ZAP-70 regulates T-cell signaling": Gangopadhyay_MannaEtal_SI

### Materials and Methods

#### Constructs:

ZAP70 tSH2 wildtype (amino acid residue number 1-256) cloned into the pSKB2 vector was gifted from Prof. John Kuriyan, UC Berkeley. The wildtype tSH2 construct of Syk (amino acid residue number 7-263) cloned into the pGEX6P1 vector was a gift from Bruce Mayer (Addgene plasmid (Syk(NC)-SH2) # 46521; http://n2t.net/addgene:46521; RRID:Addgene_46521). The point mutation in the tSH2 construct (R39A, R43P, F117A, W165C, R192A) was introduced by site-directed mutagenesis using PCR.

#### Expression and purification of tSH2 domain

The tSH2 domain of ZAP70 was expressed in *E.coli*-BL21(DE3) cells, induced with 1mM IPTG for overnight at 18°C and purified as described earlier(1). Cells were lysed by sonication or French-press in lysis buffer containing 50mM Tris, pH 8, 200mM NaCl, 20mM imidazole, 5mM β- mercaptoethanol, 5% glycerol. The tSH2 domain was purified by using Ni-NTA column and eluted in elution buffer (50mM Tris, pH 8, 200mM NaCl, 500mM imidazole, 5mM β-mercaptoethanol, 5% glycerol). The protein was further purified using a

Q-column and a gel-filtration chromatography, respectively. The purified tSH2 domain was buffered exchanged to 20mM Tris, pH 8, 150mM NaCl, 5mM β-mercaptoethanol, 5% Glycerol, concentrated to 1.5mg/ml and stored at -80°C freezer. For the NMR sample preparation, the poly(6)-His tag was removed by prescission protease digestion before purification through Q-column.

The GST-fusion tSH2 domain of Syk was expressed in *E.coli*-BL21(DE3) cells and purified as described earlier(2, 3). The cells were lysed by sonication in lysis buffer containing 1X PBS, pH 7.4 and treated with 1% TritonX100 after cell lysis. The tSH2 domain was purified using GST-column and eluted in reduced glutathione containing elution buffer (50mM Tris-HCl pH 8.0, 10mM Glutathione Reduced, 10% Glycerol). The protein was buffer exchanged to glutathione free elution buffer (50mM Tris, pH8.0, 150mM NaCl, 5mM β-mercaptoethanol, 10% Glycerol) and the GST-tag was removed by overnight digestion with prescission protease. The digested sample was further purified using GST-column and gel filtration chromatography, respectively. The purified tSH2 domain was buffer exchanged to storage buffer (50mM Tris, pH8.0, 150mM NaCl, 5mM β-mercaptoethanol, 10% Glycerol) and concentrated at 1.5mg/ml before storing at -80˚C freezer.

**Fluorescence spectroscopy:**

*Fluorescence titration:* The interaction of various tSH2 constructs and the doubly-phosphorylated ITAM-1 from CD3ζ (ITAM-Y2P-ζ1) peptide were measured by following the intrinsic tryptophan fluorescence of the protein. Syk family kinases have three conserved tryptophan residues in the tSH2 domain, W133 at the interdomain -A, W165 and W235 in the C-SH2 domain located at the interface of N-SH2 and C-SH2 domains, respectively(4) (Figure S1). The ITAM-Y2P-ζ1 was purchased from Biotechdesk®. The doubly phosphorylated ITAMs from CD3 or ζ chains bind to the tSH2 domain with hierarchical affinity(5). We choose ITAM-1 from ζ chain due to its strong (nM) binding to the tSH2 domains for our studies. Fluorescence titration was carried out in PTI Spectrofluorimeter at $\lambda_{ex}$ of 295 nm, and fluorescence emission was scanned from 300 – 400nm(6). For each titration 200µl of 0.5µM tSH2 protein was mixed with the indicated concentration of ITAM-Y2P-ζ1, given in the respective figures. The data were analyzed by fitting the curve of the $F_0/F$ *vs.* ligand concentration using the following equation:

$$F_0/F=B_{max}*X^h/(K_d^h + X^h)$$

Where $F_0$ and F is fluorescence intensity in absence and presence of ligand respectively, Bmax is the maximum binding, $K_d$ is the dissociation constant, and *h* is the Hill coefficient. The Hill coefficient ($n_H$) was also determined from the slop of Hill plot(7) from the linear fitting of the log of fraction bound *vs.* log of ligand concentration using the following equation:

$$\log\left(\frac{Y}{1-Y}\right) = n\log[L] - \log K_d$$

Where, $Y = \frac{[F-F_0]}{\Delta F_{max}}$, where F and $F_0$ are the fluorescence intensities in the presence and absence of ITAM-Y2P-ζ1, respectively. $\Delta F_{max}$ is the maximum change in fluorescence during the titration. The Hill coefficients are estimated independently from the slope of the first phase and second phase of the curve ($n_{H1}$ and $n_{H2}$, respectively)

*Fluorescence polarization assay:* The interaction between tSH2 domain and ITAM-Y2P-ζ1 was quantitatively determined from the steady-state anisotropy measurements of Alexa Fluor ™ 488 labeled ITAM-Y2P-ζ1 peptide. The labeled peptide was obtained from GenPro Biotech. The tSH2 domain and ITAM-Y2P-ζ1 interaction was measured by titrating 25nM of the labeled ITAM-Y2P-ζ1 peptide to the tSH2 proteins at indicated concentration in respective figures. The peptide and the protein solutions were prepared in 20mM Tris, pH 8, 150mM NaCl, 5mM β-mercaptoethanol, 5% Glycerol and the fluorescence measurements were recorded using Hitachi model F-4500 spectrofluorometer, equipped with a polarization accessory. Dissociation constant was determined by fitting the curve to the following equation using PRISM software(8):

$$Y=Y_0+[B_{max}*X/(K_d+X)]$$

Where Y is the measured anisotropy value in the absence of tSH2 protein, $Y_0$ is the anisotropy value of the free Alexa Fluor488 labeled ITAM-Y2P-ζ1, X is the concentration of the protein, $B_{max}$ is the maximum value of anisotropy measured and $K_d$ is the dissociation constant.

*Acrylamide quenching assay:* The conformation of tSH2 domain in the closed or open state was examined by measuring the protection of tryptophan fluorescence from acrylamide quenching. The tryptophan fluorescence for the *apo* or *holo* tSH2 domain was measured by titrating increasing concentration of acrylamide at $\lambda_{ex}$ at 295 nm. The fluorescence emission was scanned from 300-400nm, and the maximum fluorescence intensity was plotted against the acrylamide concentration. The Stern-Volmer quenching constant ($K_{sv}$) was determined by fitting the data using the following equation(9):

$$F_0/F=1+K_{sv}.X$$

Where $F_0$ is the fluorescence in absence of acrylamide, F is the measured intrinsic fluorescence; X is the concentration of the acrylamide.

**Thermal unfolding assay:**

Circular dichroism (CD) spectra were recorded on a Jasco-J800 spectropolarimeter. The wildtype and mutants tSH2 domains were buffer exchanged to 20mM phosphate buffer (20mM sodium phosphate, pH8.0, and 150mM NaCl) before recording the spectra. Two CD spectra were recorded for each construct of tSH2 protein in the *apo* state and the *holo* state (1:1 complex of tSH2: ITAM-Y2P-ζ1). Spectra were recorded from 300 to 200 nm at a temperature ranging from 20° to 60˚C with an increment 4°C. The ellipticity data was converted into molar ellipticity using the following equation.

$$[\Theta] = m^\circ * M/(10 * L * C)$$

Where [Θ] is the molar ellipticity, m° is ellipticity of the sample measured, M is average molecular weight (g/mol), L is path length of the cell (cm), and C is a concentration in g/L.m. The melting point ($T_m$), was determined from the plot of molar ellipticity at 222nm *vs.* temperature.

**Isothermal Calorimetry:**

Isothermal Titration Calorimetric studies were performed using MicroCal-autoITC (GE Healthcare) from DBT-IPLS (University of Calcutta) and Malvern-PEAQ-ITC in IISER-Kolkata. The wildtype and the mutant tSH2 domains were buffer exchanged to glycerol free buffer containing 20mM Tris, pH 8, 150mM NaCl, 5mM β-mercaptoethanol and concentrated to a concentration of 20μM. The ITAM-Y2P-ζ1 motif at a concentration of 66.6μM or 100μM was prepared in the same buffer. All the ITC titrations were measured at 20$^o$C, with nineteen injections of 2μl each and delay of 180 Sec between the injections. During titration, the protein solution was stirred at 300 rpm. The $K_a$, ΔH, and ΔS were determined from fitting the titration data using the program ORIGIN. During the curve-fitting, the data did not fit reliably to the one-site binding model or two-site independent binding model but could be fitted to the two-site sequential binding model. The ΔG (Gibb's free energy) was calculated using the equation:

$$\Delta G = \Delta H - T\Delta S$$

All the thermodynamic parameters for various tSH2 constructs binding to ITAM-Y2P-ζ1 peptide are listed in Table S6.

**NMR spectroscopy**

*NMR sample preparation:* For our NMR studies, we used the wildtype tSH2 construct (Residue number 1-256). The protein was expressed and purified, as explained above. Uniformly labeled $^{15}N$, $^{13}C$, and $^{2}H$ sample was prepared by growing the bacteria in M9 media containing $^{15}N$ ammonium chloride, $^{13}C^{2}H$ glucose, and 100% $^{2}H_2O$. The doubled labeled ($^{15}N$ and $^{13}C$), and $^{15}N$ labeled samples were prepared by growing the bacteria in M9 media containing $^{15}N$ ammonium chloride and $^{13}C$ glucose. The NMR sample of tSH2 domain was prepared by concentrating the protein to 0.35mM in sample buffer composed of 20mM Tris, pH8.0, 100mM NaCl, 1mM $Na_3VO_4$, 1mM TCEP, 0.01% Sodium azide, 10% $D_2O$.

*NMR Spectroscopy:* All NMR experiments were performed at $^{1}H$ frequencies of 900 MHz and 700 MHz on a Bruker Avance spectrometer fitted with TCI cryo-probes at 300 K, located at the University of California Berkley, USA and Bose Institute, India, respectively. Data were processed with NMRPipe(10) and analyzed using Sparky3.111(11). The 2D-$^{15}N$-$^{1}H$-HSQC and TROSY spectra were acquired with spectral width of 35 and 14 ppm for the $^{15}N$ and $^{1}H$ dimensions, respectively and with 128 ($^{15}N$) and 1024 ($^{1}H$) complex points. During the experiment, the carrier frequencies of the proton and nitrogen channels were centered at 4.7 ppm and 118 ppm, respectively. Backbone chemical shifts were assigned by using standard sets of TROSY-based triple resonance experiments: HNCO, HNCA, HN(CO)CA, CBCA(CO)NH and HNCACB(12-14). $^{15}N$ edited 3D NOESY data were acquired with 100ms mixing times. Around 85% of backbone amide resonances for the tSH2 domain in the *holo* sate were assigned.

*NMR Titration:* The interaction between the tSH2 domain of ZAP-70 and ITAM-Y2P-ζ1 peptide was determined from the NMR titration. Several $^{15}N$-$^{1}H$ HSQC spectra of tSH2 domain were recorded at different protein to peptide ratio. The NMR titration was followed by measuring the $^{15}N$-$^{1}H$ HSQC spectra of 1:2 complex of 15N labeled tSH2 domain and unlabeled ITAM-Y2P-ζ1 peptide. Subsequent samples of 1:1, 1:0.8, 1:0.6, 1:0.4 and 1:0.2 complex of tSH2 domain and ITAM-Y2P-ζ1, respectively, were prepared by serially diluting the tSH2:ITAM-Y2P-ζ1 complex with $^{15}N$ labeled tSH2-*apo* protein. The backbone chemical shift assignments were transferred by comparing the assignment of 1:1 complex of tSH2: ITAM-Y2P-ζ1. The titration data were analyzed from the chemical shift difference and the peak intensity measured from for each titration point. The compounded $^{1}H$, $^{15}N$ chemical shift differences (□CCS) between 1:2 complex of tSH2: ITAM-Y2P-ζ1 and subsequent dilution steps were computed by using $[((\delta^{1}_{HN})^2+(\delta^{15}_{NH}/6.5)^2)^{1/2}]$[15]. To find out the chemical shift changes of the amide backbone on binding to ITAM-Y2P-ζ1, we measure the □□CCS. The □□CCS chemical shifts were determined from the difference between the □CCS for 1:2/ 1:1 tSH2: ITAM-Y2P-ζ1 complex and the □CCS for 1:2/1:0.8, 1:2/1:0.6 or 1:2/1:0.4 complex. The peak intensity was determined from the cross-peak fit height by using the Gaussian line fitting protocol implemented in Sparky 3.111. The normalized peak intensity ($I/I_0$) was determined from the ratio of peak intensity measured at respective protein to peptide ratio to that of 1:2 complex of tSH2: ITAM-Y2P-ζ1. The predicted Cα chemical shifts from the crystal structure of tSH2 domain in complex with doubly phosphorylated ITAM (PDB: 2OQ1) was derived by using SHIFTX2(16).

**Residue Network Analysis**

The two-dimensional residue interaction network (RIN) for the *apo* and *holo* tSH2 domain of ZAP70 was constructed from the PDB structure 1M61 and 2OQ1, respectively. The undirected RIN was generated from the three-dimensional structure of each protein using RINERATOR module of RINalyzer(17, 18), and Cytoscape was used for the network analysis and visualization(19, 20). In the RIN, each node represents the amino acid residue in the protein, and the edges are noncovalent interactions (namely interatomic contact, hydrogen bond, and overlapping Van der Waals radii) between the side-chains of the amino acids that link two nodes in the protein structure. Any two amino acid residues i and j are considered to be in contact if the distance between i and j is less than 5.0 Å, which approximates the upper limit for attractive London–van-der-Waals forces(21, 22). The hydrogen atoms were added to the protein structure in stereochemically favorable positions by using the program REDUCE(23), and the noncovalent interactions between two residues were identified using the program PROBE(24). After construction of the interaction network, the Network Analyzer(25) plugin of Cytoscape was used to calculate the topological parameters like degree, clustering coefficient (C), shortest path length (L) and the average number of neighbors (<N>). Random networks of tSH2 domain of ZAP70 in *apo* and complex with ITAM-Y2P was created using network randomizer. The predicted average clustering coefficient ($C_R$) and average shortest path length ($L_R$) were used to evaluate the properties of the network(26). The shortest intermolecular undirected network coupling the phosphate-binding pocket of C-SH2 domain to the N-SH2 domains was determined by tracing the path connecting the immediate neighbors of phosphotyrosine at the C-SH2 domain to the first neighbors of phosphotyrosine at N-SH2 domain. For easier identification of hubs, the nodes having a high degree was assigned a higher height and width.

**Molecular Dynamics Simulation:**

*System Preparation and Simulation Details:* Molecular dynamics (MD) simulation was carried out on eight different systems (S1 to S8) as given in Table S2. Two systems with the tSH2 domain of wild type ZAP-70 protein were prepared for simulation, including tSH2-*holo*(S1) and tSH2-*apo* (S2) structures. The crystal structures of the doubly phosphorylated ITAM bound tSH2 domain (tSH2-*holo*) of ZAP-70 (PDB ID: 2OQ1), and tSH2-*apo* (PDB-ID:1M61) were obtained from the Protein Data Bank(27, 28). To understand the domain-specific interactions of N-SH2 and C-SH2 domain with phosphorylated ITAM, two ITAM modified structures were prepared (Figure 2a). In the first structure (named: N-SH2$^{ITAM\text{-}YP}$), N-SH2 phosphate-binding pocket was occupied by the phosphotyrosine residue of ITAM, and the C-SH2 phosphate-binding pocket was unoccupied. In the second structure (named: C-SH2$^{ITAM\text{-}YP}$) the N-SH2 phosphate-binding pocket was unoccupied and the C-SH2 bound to the phosphotyrosine (Figure 2A). In case of N-SH2$^{ITAM\text{-}YP}$ (S3) and C-SH2$^{ITAM\text{-}YP}$ (S4) systems, the residues 12 to 19 and residues1 to 12 of the doubly phosphorylated ITAM were retained in the structure, respectively. In addition, four in-silico mutants at the allosteric network region were made from the tSH2-*holo* (PDB ID: 2OQ1) structure i.e. tSH2-*holo*$^{W165C}$ (S5), tSH2-*holo*$^{F117A}$ (S6), tSH2-*holo*$^{R43A}$ (S7) and tSH2-*holo*$^{R43P}$ (S8). Each construct was solvated using the TIP3P water model(29) in orthogonal simulation box with 12Å padding of water. Sodium chloride (NaCl) was added to adjust the salt concentration to 150mM in each system. The AMBER-ff14SB force field was applied for the protein, and all simulations were carried out using AMBER 18(30) under periodic boundary condition. The temperature was kept constant at 300K by applying Langevin thermostat. Particle-mesh-Ewald method(31) was used for the calculation of long-range electrostatics with an interaction cut-off of 10Å. Bonds involving hydrogen atoms were constrained using the SHAKE algorithm(32). Energy minimization was carried out for 10000 steps, followed by a system equilibration phase for 1 ns. MD simulation was performed with a time integration step of 2 fs under NPT ensemble for 200 ns for S1 and S2 whereas 100 ns for the rest of the systems (S3 to S8). Trajectory data was saved at every 1 ps interval for analysis. Simulation trajectories were analyzed by CPPTRAJ(33).

*Interaction Energy (IE) Analysis:* The energetics of the doubly-phosphorylated ITAM peptide and tSH2 domain of ZAP-70 interaction was evaluated from average non-bonded interaction energy (in kcal/mol) determined throughout the simulation trajectories in respective systems. The interaction energy comprises of both the electrostatic and Van der Waals components and calculated for all atom pairs between the doubly-phosphorylated ITAM peptide and the tSH2 domain. To study the difference of interaction strength between the phosphotyrosine motif of ITAM and two SH2 domains of the ZAP-70, interaction energy was calculated for phosphotyrosine residue (residue no. 15) - N-SH2 domain (residue no. 1 to 110) and phosphotyrosine residue (residue no. 4) – C-SH2 domain (residue no. 163 to 254), respectively. To compare the interaction energy in the *holo* and mutated structures of tSH2 domain the entire doubly phosphorylated ITAM (Res 1 to 19) and the tSH2 domain of ZAP-70 (Res 1 to 254) were considered.

**Table S1: Reported dissociation constant forITAM-Y2P-ζ1 and tSH2 domain of ZAP-70**

| Name of the protein | $K_d$ | Method | Reference |
|---|---|---|---|
| [a]ZAP-70 tSH2 | 56.7nM | Surface Plasmon Resonance | Bu et. al., 1995, PNAS(34) |
| [a]ZAP-N*C (R37K) | 3.5μM | Surface Plasmon Resonance | Bu et. al., 1995, PNAS(34) |
| [a]ZAP-NC*(R190K) | 1.8μM | Surface Plasmon Resonance | Bu et. al., 1995, PNAS(34) |
| [a]Zap-70 tSH2 | 45.2nM | Surface Plasmon Resonance | Labadia et. al., 1996, Journal of Leukocyte Biology(35) |
| [a]ZAP-70 tSH2 | 3.5nM | Surface Plasmon Resonance | Labadia et. al., 1996, Journal of Leukocyte Biology(35) |
| [a]ZAP-70 tSH2 | 3.5nM | Surface Plasmon Resonance | Labadia et. al., 1997, Archives of Biochemistry and Biophysics(5) |
| [a]ZAP-70 N-SH2 (R187K) | 47.6μM | Surface Plasmon Resonance | Labadia et. al., 1997, Archives of Biochemistry and Biophysics(5) |
| [a]ZAP-70 C-SH2(R43K) | 17.8μM | Surface Plasmon Resonance | Labadia et. al., 1997, Archives of Biochemistry and Biophysics(5) |
| [a]Zap-70 tSH2 | 2nM | Surface Plasmon Resonance | Ottinger E.A., et. al., 1998, Journal of Biological Chemistry(36) |
| [a]ZAP-70 tSH2 | 33nM | Isothermal Titration Calorimetry | O'brien et. al., 2000 Protein Science(37) |
| [b]ZAP-70 full-length | 76.6nM | Fluorescence Polarisation | Deindl S., et. al., 2009, PNAS(38) |
| [b]ZAP-70 full-length W131A | 15.2nM | Fluorescence Polarisation | Deindl S., et. al., 2009, PNAS(38) |
| [b]ZAP-70 full-length YYFF | 96.1nM | Fluorescence Polarisation | Deindl S., et. al., 2009, PNAS(38) |

[a] ZAP70 tSH2 domain was used in these experiments.[b] ZAP70 full-length constructs were used in these experiments.

**Table S2: Details of Molecular Dynamics simulation studies**

| Sl No. | System | System Construct(s) | Simulation Time (ns) | Remark |
|---|---|---|---|---|
| 1 | S1 | tSH2-*holo* (PDB ID: 2OQ1) | 200 | Wild- type |
| 2 | S2 | tSH2-*apo* (PDB ID: 1M61) | 200 | |
| 4 | S3 | N-SH2$^{ITAM-YP}$ (Phosphate binding pocket) | 100 | ITAM-Y2P Modified |
| 5 | S4 | C-SH2$^{ITAM-YP}$ (Phosphate binding pocket) | 100 | |
| 6 | S5 | tSH2-*holo*$^{W165C}$ | 100 | Mutant |
| 7 | S6 | tSH2-*holo*$^{F117A}$ | 100 | |
| 8 | S7 | tSH2-*holo*$^{R43A}$ | 100 | |
| 9 | S8 | tSH2-*holo*$^{R43P}$ | 100 | |

**Table S3: Inter-residue distances of the networking residues**

| Systems | Distance (Å) | | | | | | | | |
|---|---|---|---|---|---|---|---|---|---|
| | R43(O)-Q236(NE2) | F117(CG)-Q236(CG) | W235(CH2)-Q236(NE2) | F117(CD2)-W235(CZ3) | W165(CZ2)-W235(CB) | W165(CB)-L190(CB) | L190(CD1) - R192(CG) | W165(CZ3)-L191(CD1) | L191(CD2)-R192(O) |
| tSH2-*holo* (S1) | 4.63± 0.59 | 4.58± 0.54 | 4.41± 0.54 | 5.24± 0.83 | 4.94± 1.03 | 5.50± 0.37 | 7.42± 0.92 | 4.93±0.70 | 5.17± 0.74 |
| tSH2-*holo*$^{W165C}$ (S6) | 2.99± 0.34 | 5.63± 0.48 | 12.63± 0.99 | 15.71± 0.75 | - | 10.99± 1.71 | 6.10± 0.62 | - | 6.75± 0.67 |
| tSH2-*holo*$^{F117A}$ (S7) | 5.48± 1.63 | - | 7.22± 0.93 | - | 8.52± 1.00 | 5.31± 0.76 | 5.56± 0.96 | 6.62±0.96 | 6.11± 1.09 |
| tSH2-*holo*$^{R43A}$ (S8) | 7.28± 1.70 | 11.70± 1.82 | 6.91± 1.81 | 8.46± 2.69 | 12.15± 1.32 | 6.07± 0.64 | 6.99± 0.99 | 9.61± 1.30 | 6.14± 0.54 |
| tSH2-*holo*$^{R43P}$ (S9) | 4.78± 1.17 | 6.62± 1.04 | 7.07± 2.01 | 6.58± 1.90 | 6.37± 1.43 | 8.94± 1.34 | 5.86± 0.76 | 6.32± 1.13 | 7.82± 0.63 |

**Table S4: Comparison of residue interaction networks between the *apo* and ITAM-Y2P-ζ1 bound state of tSH2 domain of ZAP-70**

| RIN | Number of nodes in RIN | Number of edges in RIN | <N> | C | $C_R$ | $C/C_R$ | L | $L_R$ | $L/L_R$ |
|---|---|---|---|---|---|---|---|---|---|
| tSH2-*apo* ZAP-70 (PDB ID: 1M61) | 259 | 2630 | 7.344 | 0.445 | 0.079 | 5.633 | 5.385 | 2.108 | 2.555 |
| ITAM ZAP-70 (PDB ID: 2OQ1) | 273 | 3305 | 7.612 | 0.465 | 0.087 | 5.344 | 5.272 | 2.017 | 2.614 |

RIN, residue interaction networks of different protein structures; Number of nodes in RIN, amino acid residues in the protein; Number of edges in the RIN; <N>, average number of neighbors; C, average clustering coefficient; $C_R$-average clustering coefficient for the random networks with the same size; $C/C_R$, average clustering coefficient ratio(26); L, average shortest path length; $L_R$-average shortest path length for the random networks with the same size; $L/L_R$, average shortest path length ratio(26). A high $C/C_R$ ratio indicates that the residues of the network are highly clustered.

**Table S5: Thermodynamic parameters forZAP-70 tSH2 domain binding to ITAM-Y2P-ζ1**

| Construct | ΔG1 (cal/μmol) | ΔH1 (cal/μmol) | TΔS1 (cal/μmol) | ΔG2 (cal/μmol) | ΔH2 (cal/μmol) | TΔS2 (cal/μmol) |
|---|---|---|---|---|---|---|
| tSH2 wildtype | -447.2 ± 187.2 | -480.7 ± 176.9 | -33.5 ± 12.1 | 443.3 ± 1.8 | 478.7 ± 1.9 | 35.4 ± 13.2 |
| tSH2 F117A | -178.8 ± 38.0 | -191.5 ± 40.8 | -12.8 ± 2.8 | 191.3 ± 42.6 | 205.9 ± 45.8 | 14.6 ± 3.1 |
| tSH2 W165C | -175.9 ± 51.9 | -188.9 ± 56.6 | -13.0 ± 4.7 | 182.9 ± 48.0 | 197.2 ± 52.1 | 14.3 ± 4.0 |
| tSH2 R43P | -- | -- | -- | -- | -- | -- |

ITC experiments of wildtype and mutants of ZAP-70 tSH2 domain were carried out with ITAM-Y2P-ζ1. The ΔG (cal/μmol), ΔH (cal/μmol) and TΔS (cal/μmol) are averages ± SD from 3 independent experiments.

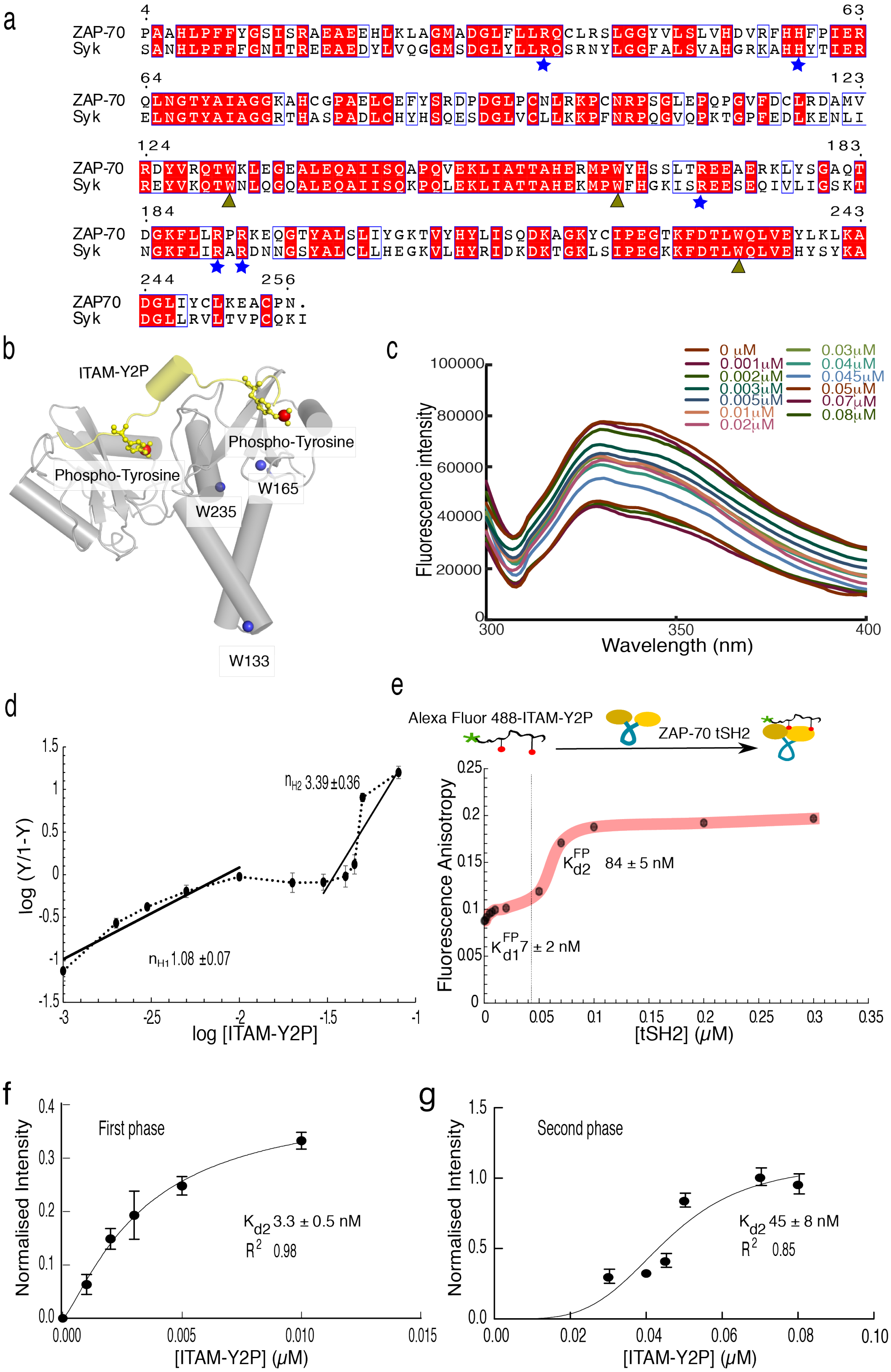


**Figure S1**: **Binding of ZAP-70 tSH2 domain and doubly-phosphorylated ITAM-Y2P-ζ1 from fluorescence measurement (related to Figure 1)** (a) Sequence alignment of tSH2 domain of Syk and

ZAP-70. The sequence similarity was 95%, the amino acid residues at the phosphate-binding pocket are indicated by star. The conserved tryptophan residues used as a probe in the fluorescence experiments are indicated by arrow-head. (b) The conserved tryptophan residues are mapped on to the tSH2-*holo* structure of ZAP-70 (PDB ID: 2OQ1). (c) Representative fluorescence intensity scan ($\lambda_{em}$ 300nm to 400nm) from the titration of tSH2 domain of ZAP-70 and ITAM-Y2P-ζ1 peptide. (d) Hill plot for the titration of tSH2 domain of ZAP-70 and ITAM-Y2P-ζ1 peptide, described in figure 1c. The Hill-coefficient ($n_H$) was determined from the slope of the plot. (e) Biphasic binding curve of the Alexa Fluor 488-ITAM-Y2P-ζ1 peptide to wildtype tSH2 domain was probed from the plot of fluorescence anisotropy *vs* tSH2 domain concentration. (f) and (g) The first phase and the second phase of binding observed during the titration of ITAM-Y2P-ζ1 to tSH2 domain (in figure 1c) was fitted separately to one-site specific binding model implemented in Prism.

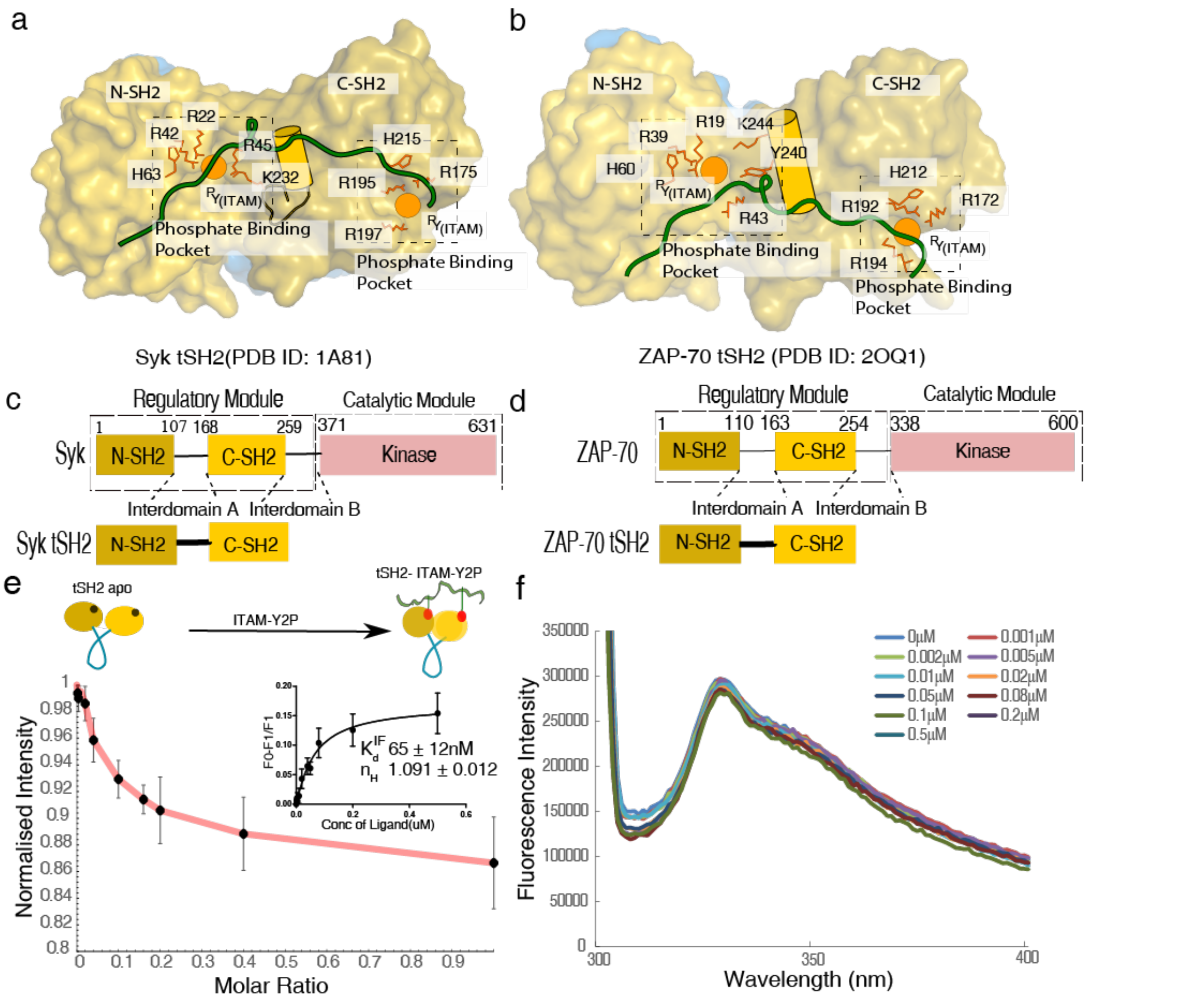


**Figure S2: Binding of Syk tSH2 domain and doubly-phosphorylated ITAM-Y2P-ζ1 peptide (Related to Figure 1)** (a) tSH2-*holo* structure of (a) Syk (PDB ID: 1A81) and (b) ZAP-70 (PDB ID: 2OQ1) are shown as space-filled model. The amino acid residues noncovalently interacting to the phosphotyrosine residue (orange circle) of ITAM-Y2P-ζ1 at the N-SH2 and C-SH2 phosphate-binding pockets are labeled. (c) Schematic representation of domain architecture of full-length Syk and tSH2 domain used in this study. (d) Schematic representation of domain architecture of full-length ZAP-70 and tSH2 domain used in this study. (e) Titration of ITAM-Y2P-ζ1 and tSH2 domain of Syk determined from the measurement of intrinsic tryptophan-fluorescence at the indicated ligand to protein molar ratio. Inset showing curve fitting for ITAM-Y2P-ζ1 binding. The error bar represents the standard deviation from three experiments. (f) Representative fluorescence intensity scanned ($\lambda_{em}$ 300nm to 400nm) from the titration of tSH2 domain of Syk and ITAM-Y2P-ζ1 peptide.

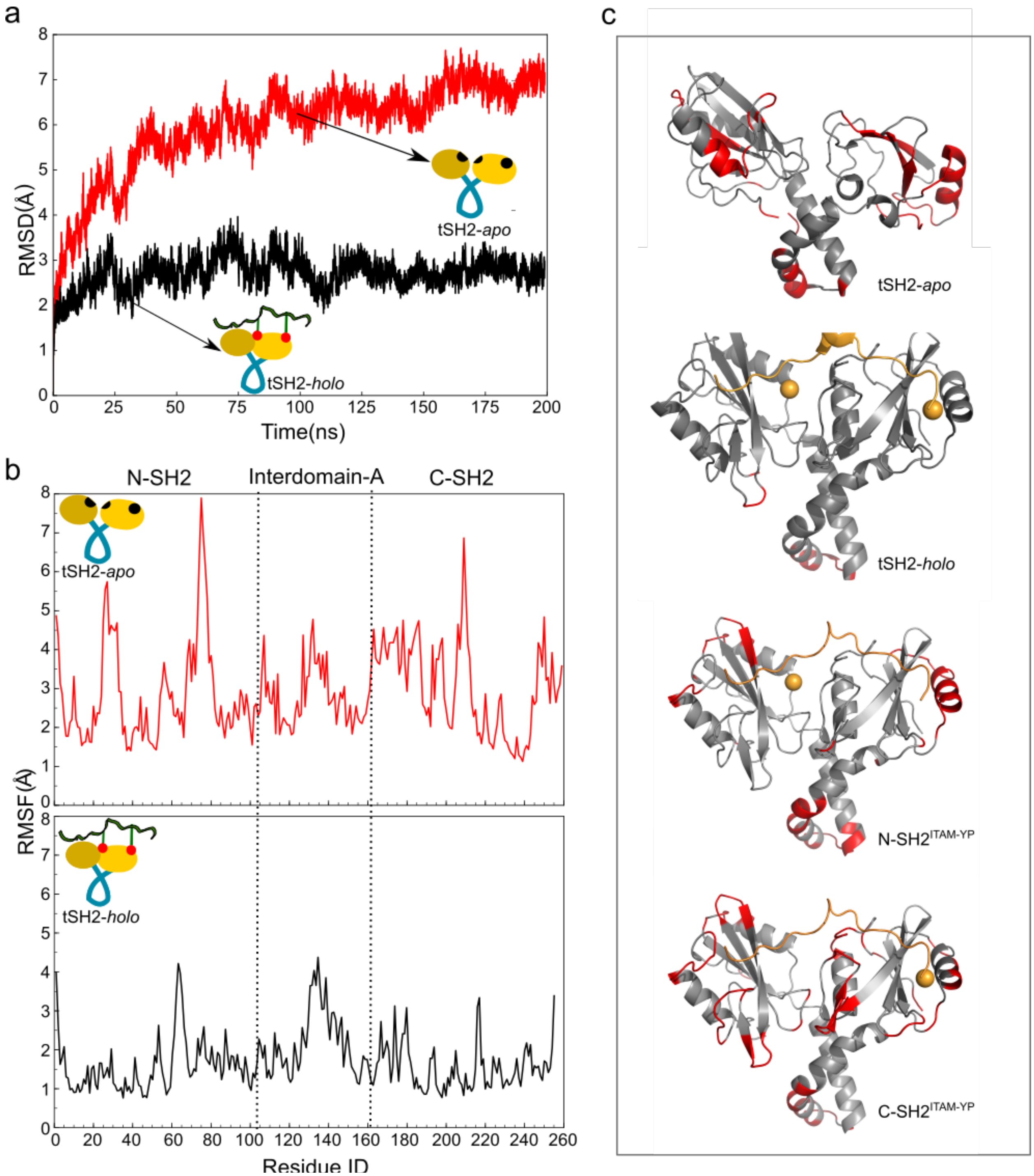


**Figure S3: Structural evolution of the tSH2-*holo* and tSH2-*apo* structures during MD simulation (Related to Figure 2)** (a) Cα root-mean-squared deviation (RMSD) of the tSH2-*holo* and tSH2-apo structure of ZAP-70 observed throughout the 200 ns simulation trajectory. (b) Average root-mean-square fluctuation (RMSF) is plotted against the residue number. (c) Residue specific RMSF greater than 3.5 Å from respective molecular dynamics simulations are mapped on to the tSH2-*holo* structure.

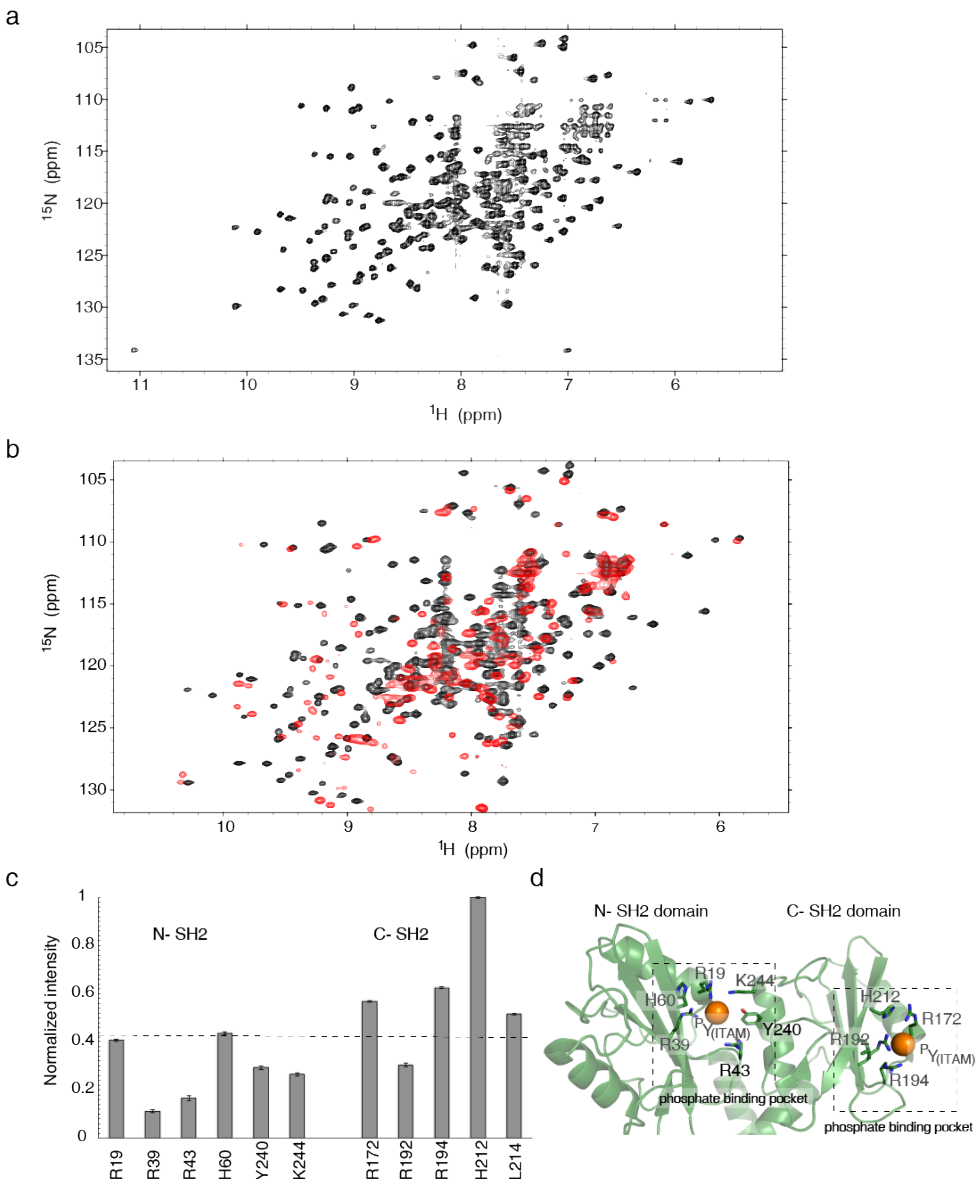

a
$^{15}$N (ppm)
$^{1}$H (ppm)
b
$^{15}$N (ppm)
$^{1}$H (ppm)
c
Normalized intensity
N- SH2
C- SH2
R19
R39
R43
H60
Y240
K244
R172
R192
R194
H212
L214
d
N- SH2 domain
C- SH2 domain
R19
K244
H60
$^{P}$Y(ITAM)
Y240
R39
R43
phosphate binding pocket
H212
R172
R192
R194
phosphate binding pocket


e

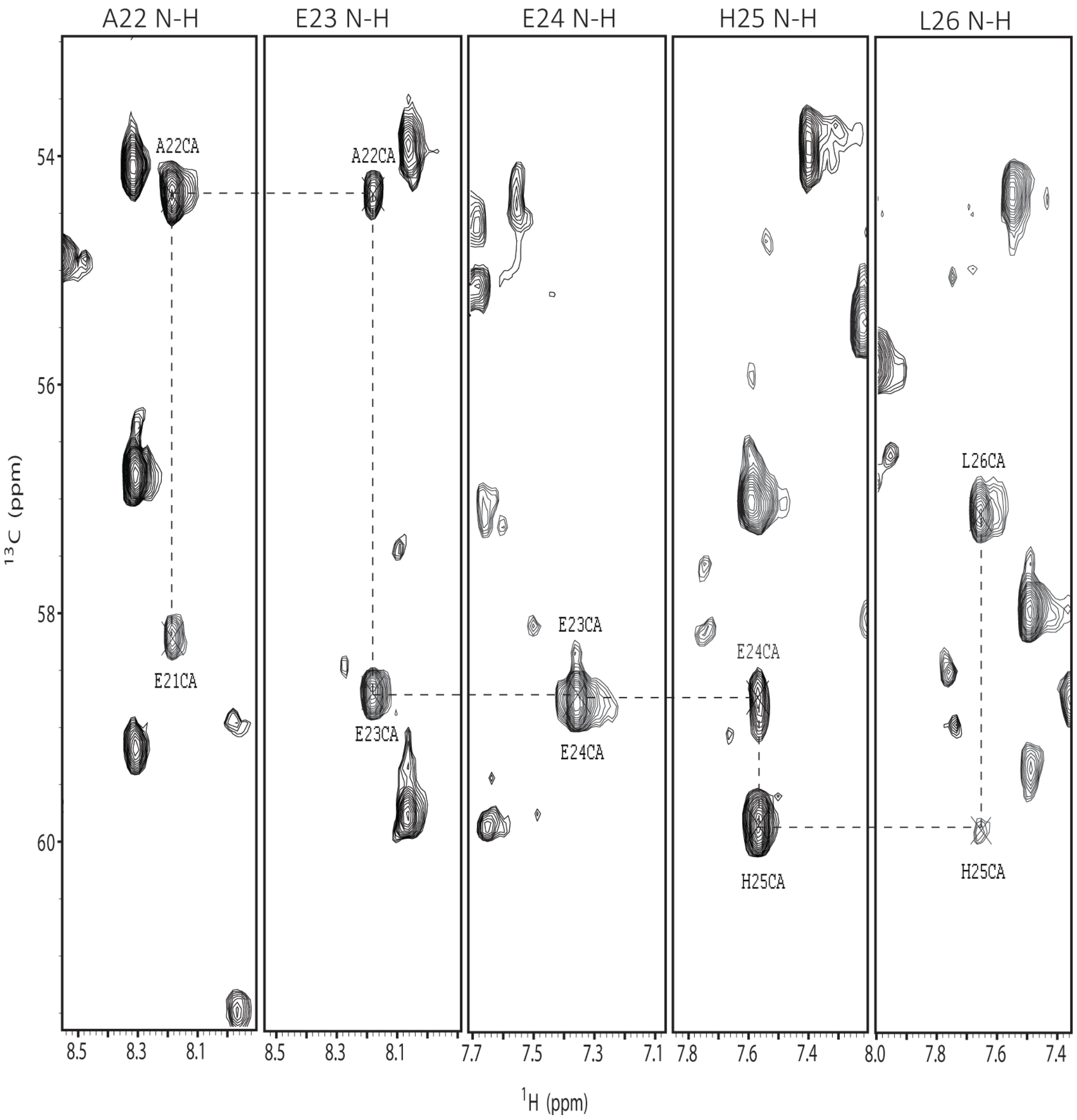


**Figure S4: (Related to Figure 3)** (a) $^{15}N$-$^{1}H$ TROSY spectra of tSH2-*holo* structure recorded at $^{1}H$ frequency of 900 MHz. (b) $^{15}N$-$^{1}H$ TROSY spectra showing the overlap of tSH2-*holo* (black) and tSH2-*apo* (red) structure. (c) Normalized intensity from the 3D HNCO spectra for the amino acid residues at the N-SH2 and C-SH2 phosphate binding pocket. The intensity for all amino acid residues plotted is normalized against the intensity of H212. The dashed line indicates the value for the average normalized intensity. (d) Represents the amino acid sizes interacting within tyrosine phosphate at the N-SH2 and C-SH2 phosphate-binding pocket. (e) Representative strip plot from HNCA experiment showing backbone assignment of tSH2 domain bound to ITAM-Y2P-ζ1.

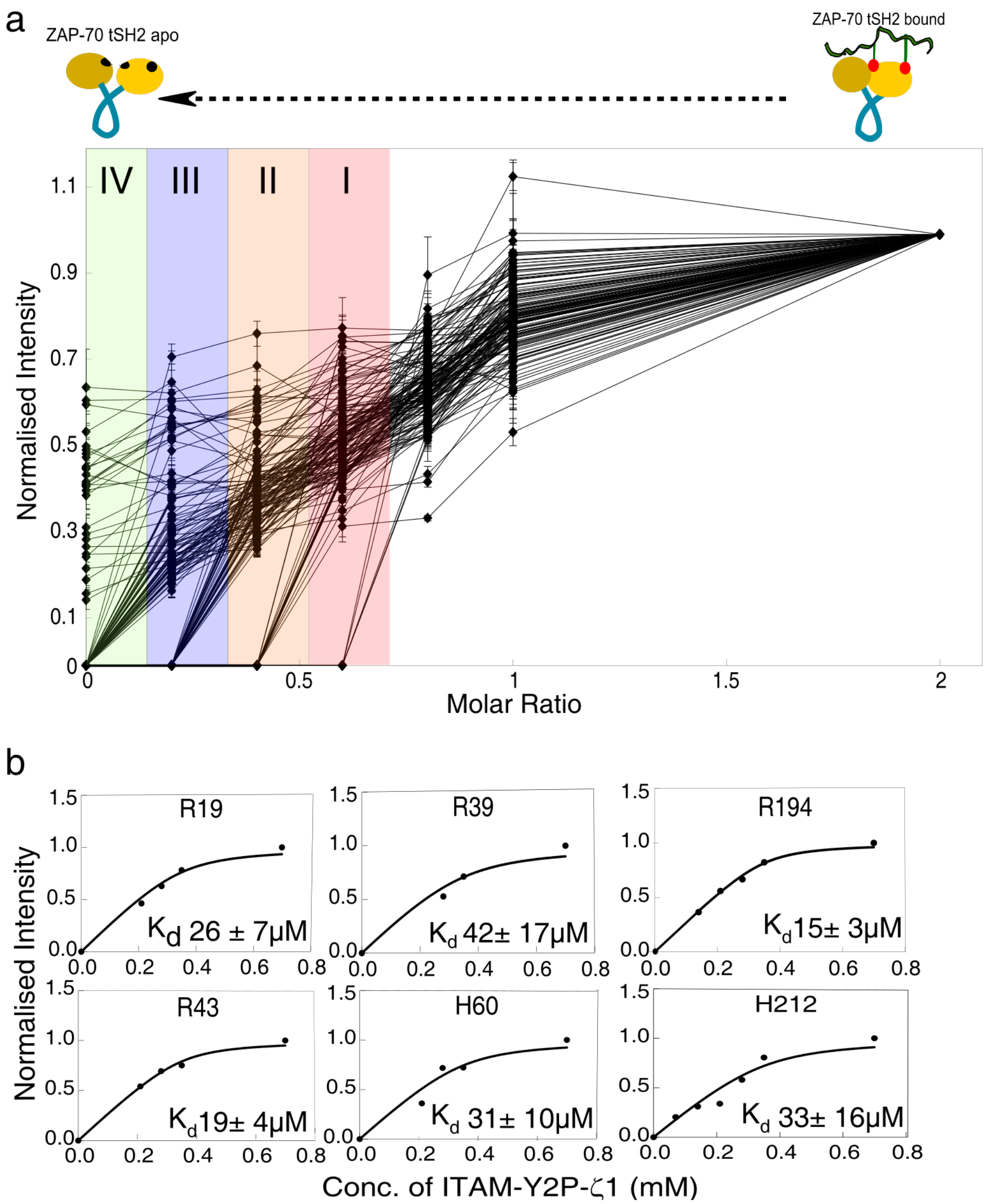


**Figure S5: NMR titration experiment of tSH2 wildtype and ITAM-Y2P-ζ1 (Related to Figure 3)** (a) Normalized intensity of backbone amide is plotted against the ligand to protein molar ratio. During the titration, tSH2-*apo* protein was added to the tSH2-*holo* sample to achieve the NMR sample at the desired molar ratio. The vertical color represents the four class of amino acid residues described in the text. The amino acid residues that line broadens beyond detection in the sample at a ligand to protein molar ratio of 0.6, 0.4 and 0.2 are grouped class-I (red), class-II (orange), and class-III (blue), respectively. All other residues are grouped into class-IV (green). The lines connecting the points are for guiding eyes. (b) The residue-specific binding constant was estimated by fitting the normalized intensity from NMR titration experiment.

a

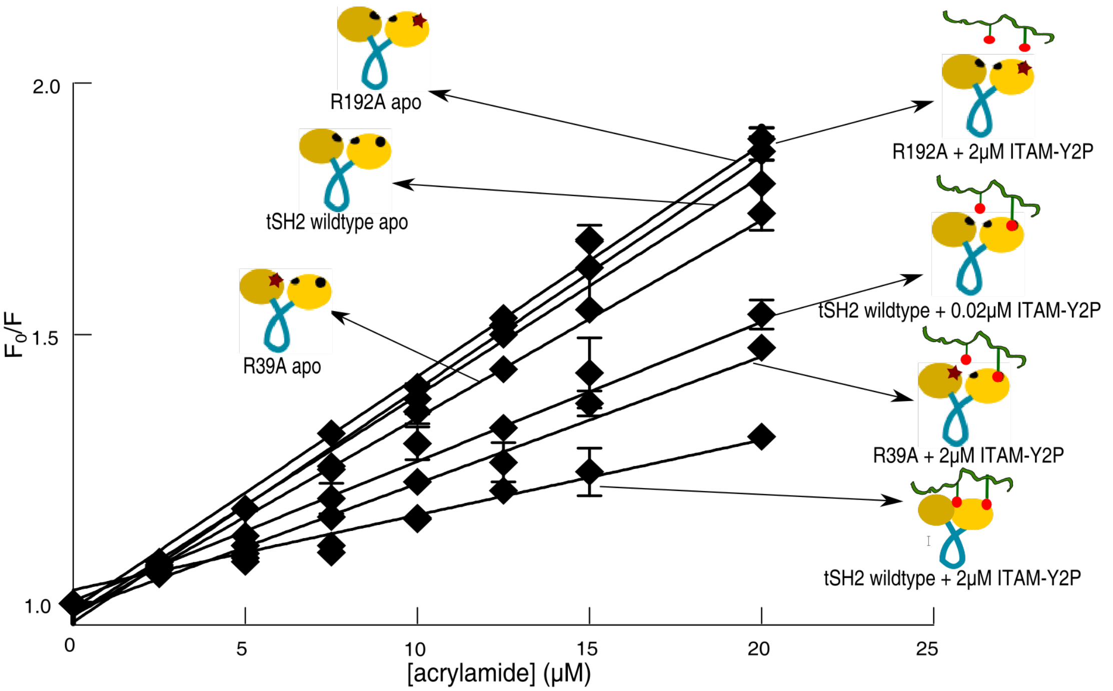


b

| Name of the protein | $K_{sv}$ ($\mu M^{-1}$) | Schematic Representation |
|---|---|---|
| tSH2 wildtype apo | 0.044 ± 0.003 | 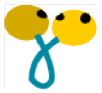 |
| tSH2 wildtype + 0.02µM ITAM-Y2P (1:0.01) | 0.027 ± 0.002 | 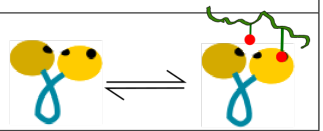 |
| tSH2 wildtype + 2µM ITAM-Y2P (1:1) | 0.015 ± 0.001 | 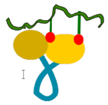 |
| R39A apo | 0.039 ± 0.003 | 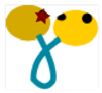 |
| R39A + 2µM ITAM-Y2P | 0.025 ± 0.003 | 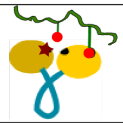 |
| R192A apo | 0.046 ± 0.004 | 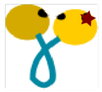 |
| R192A + 2µM ITAM-Y2P | 0.046 ± 0.002 | 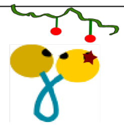 |

**Figure S6: Determination of the open or closed conformation of tSH2 domain by acrylamide quenching of tryptophan fluorescence (Related to Figure 3)** (a) The Stern-Volmer plot of normalized fluorescence intensity against increasing acrylamide concentration. The Stern-Volmer coefficient (*$K_{sv}$*) was determined from the slope of the curve. (b) Schematic representation of tSH2 samples used in this study along with the respective Stern-Volmer coefficient ($K_{sv}$). The location of the mutation is indicated by star.

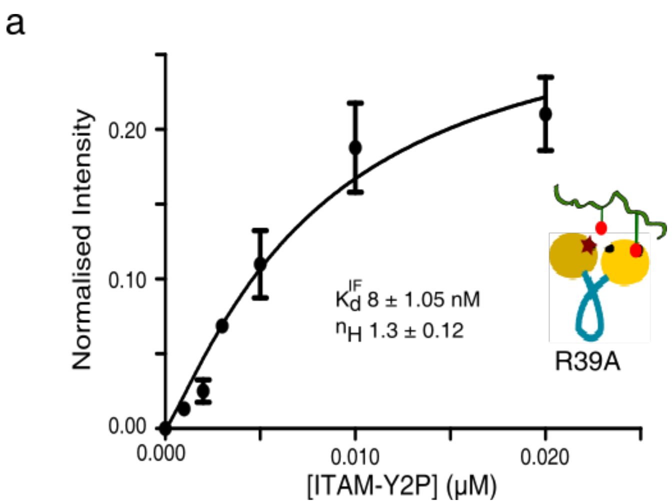


**Figure S7: (Related to Figure 3)** The change in intrinsic fluorescence for R39A mutant of tSH2 domain with increasing concentration of ITAM-Y2P-ζ1 (Figure 3) was fitted to first-order binding equation implemented in the program Prism.

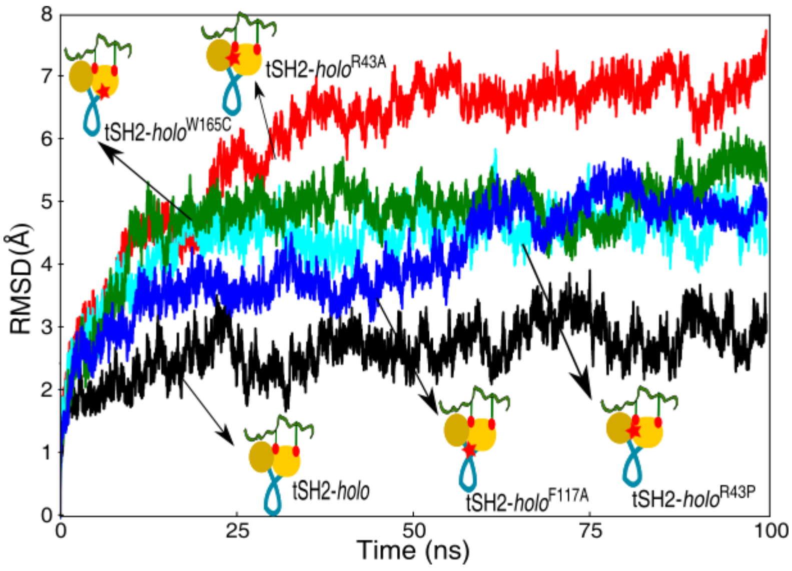


**Figure S8: Root-mean-squared deviation (RMSD) of the wildtype and four in-silico mutants in the tSH2-*holo* state during MD simulation: (Related to Figure 6)** Cα RMSD values of the tSH2-*holo*, tSH2-*holo*$^{W165C}$, tSH2-*holo*$^{F117A}$, tSH2-*holo*$^{R43A}$ and tSH2-*holo*$^{R43P}$ mutants are plotted against the simulation time. Increase in RMSD values for the mutants than the wildtype tSH2-*holo* structure suggest that the conformation of the mutated tSH2- domain structures deviated from the wildtype tSH2-holo structure.

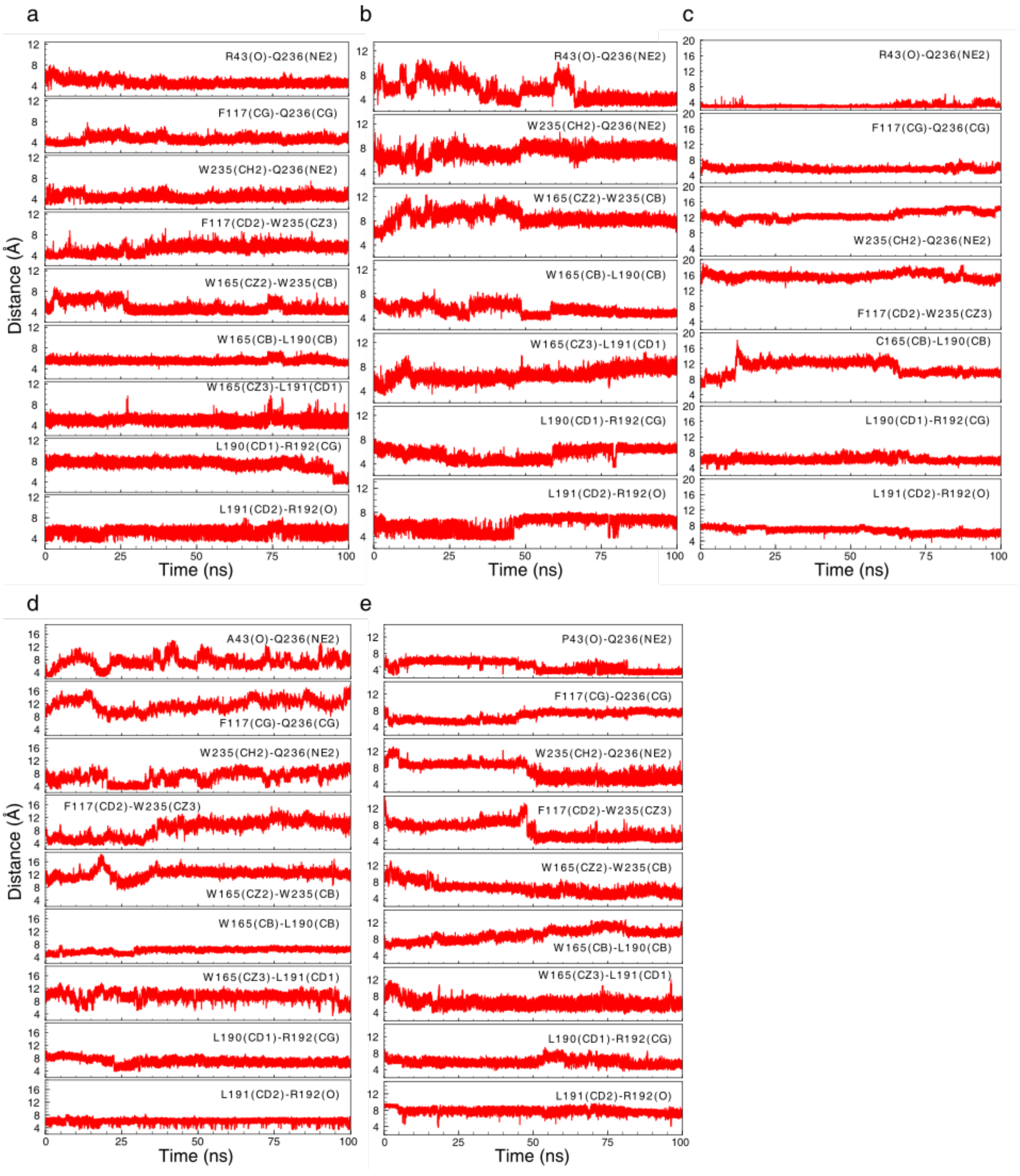


**Figure S9: Dynamic behavior of the allosteric-network during MD simulation (Related to Figure 6)**
Time evolution of the inter-residue distances between amino acid residues involved in the allosteric-network for (a) tSH2-*holo*,(b) tSH2-*holo*$^{F117A}$(c) tSH2-*holo*$^{W165C}$(d) tSH2-*holo*$^{R43A}$ and (e) tSH2-*holo*$^{R43P}$. The network is retained in the wildtype tSH2-*holo* structure, whereas most of the interactions are broken in the mutated structures.

a

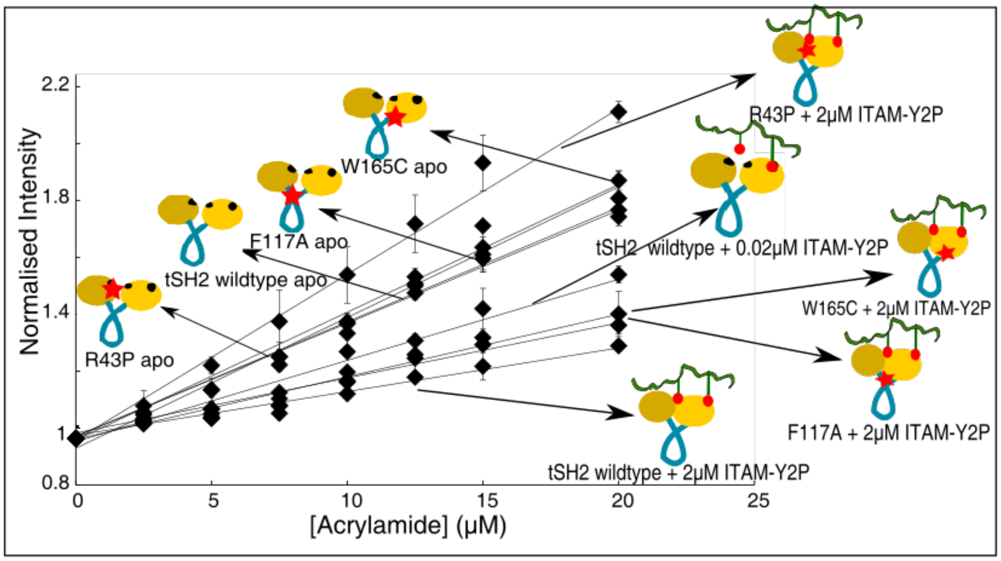


b

| Name of the protein | $K_{SV}$ ($\mu M^{-1}$) | Schematic Representation |
|---|---|---|
| tSH2 wildtype apo | 0.044 ± 0.003 | |
| tSH2 wildtype + 0.02µM ITAM-Y2P (1:0.01) | 0.027 ± 0.002 | |
| tSH2 wildtype + 2µM ITAM-Y2P (1:1) | 0.015 ± 0.001 | |
| F117A apo | 0.039 ± 0.008 | |
| F117A + 2µM ITAM-Y2P | 0.019 ± 0.004 | |
| W165C apo | 0.046 ± 0.004 | |
| W165C + 2µM ITAM-Y2P | 0.026 ± 0.01 | |
| R43P apo | 0.039 ± 0.006 | |
| R43P + 2mM ITAM-Y2P | 0.033 ± 0.004 | |

c

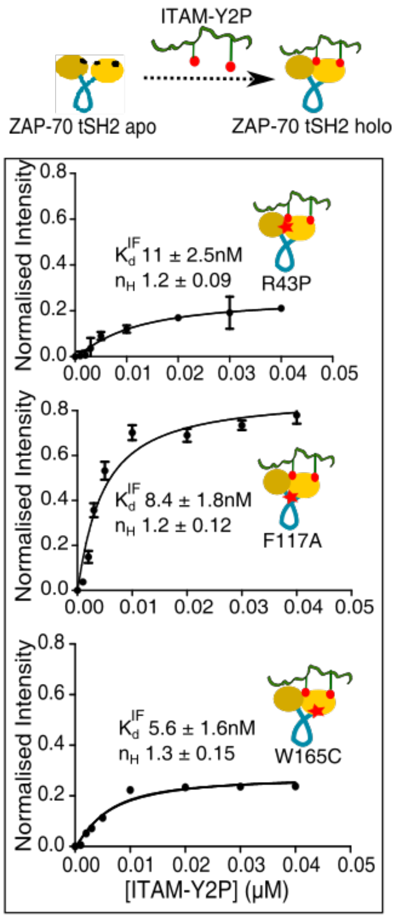


**Figure S10: Determination of the open or closed conformation of allosteric-network mutants of tSH2 domain by acrylamide quenching of tryptophan fluorescence (Related to Figure 7)** (a) The Stern-Volmer plot of normalized fluorescence intensity of wildtype, allosteric-network mutants tSH2 domain of

ZAP-70 in the *apo* and *holo* state, respectively, against increasing acrylamide concentration. The Stern-Volmer coefficient ($K_{sv}$) was determined from the slope of the curve. (b) Schematic representation of tSH2 samples used in this study along with the respective Stern-Volmer coefficient ($K_{sv}$). The location of the mutation is indicated by star. (c) The dissociation constant ($K_d^{IF}$) and Hill-coefficient ($n_H$) for the allosteric-network mutants (R43P, F117A, and W165C) was determined from the intrinsic fluorescence experiments in figure 7a by fitting onto one-site binding model implemented in the program prism.

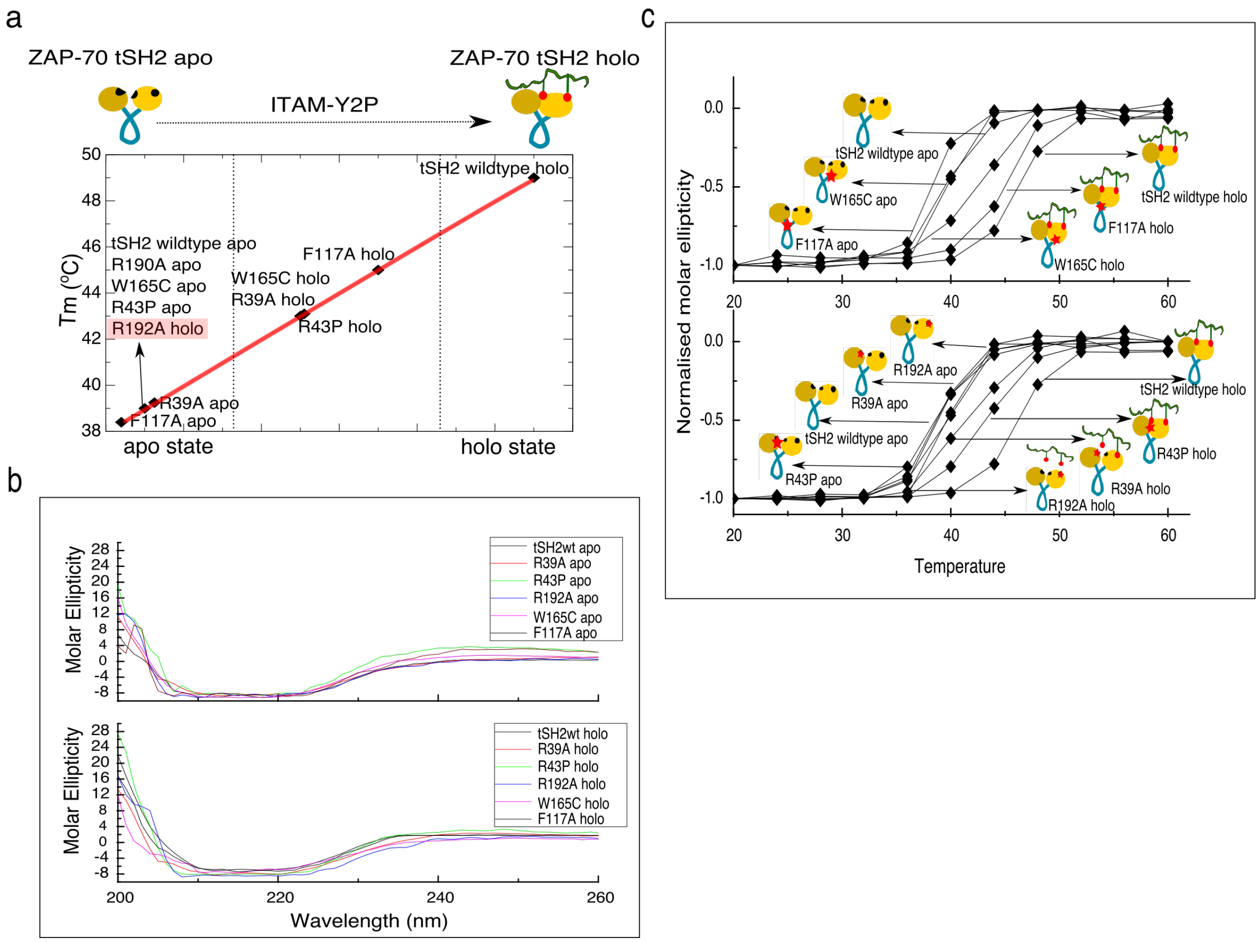


**Figure S11: Effect of tSH2 domain mutation on the thermal stability of SH2 domains upon ITAM-Y2P-ζ1 binding (Related to Figure 7)** (a) Comparative analysis of thermal denaturation profile ($T_m$) of *apo* and *holo* tSH2 domain of wildtype and the mutants. (b) Representative circular dichroism (CD) spectra of tSH2 domain of ZAP-70 wildtype, and mutants (F117A, W165C, R39A, R192A and R43P) in the *apo* (top panel) and *holo* (bottom panel) state was recorded at 20˚C. (c) The thermal unfolding of tSH2 domain of ZAP-70 wildtype, and mutants (F117A, W165C, R39A, R192A and R43P) in the *apo* (top panel) and *holo* (bottom panel) state respectively, were determined from the respective CD spectra recorded as a function of temperature ranging from 20˚C to 60˚C with an interval of 4ºC. The $T_m$ was determined from the plot of normalized molar ellipticity calculated at 222nm wavelength against the temperature.
